# Discovery of iridoid cyclase completes the iridoid pathway in asterids

**DOI:** 10.1101/2025.06.12.659277

**Authors:** Maite Colinas, Chloée Tymen, Joshua C. Wood, Anja David, Jens Wurlitzer, Clara Morweiser, Klaus Gase, Ryan M. Alam, Gabriel R. Titchiner, John P. Hamilton, Sarah Heinicke, Ron P. Dirks, Adriana A. Lopes, Lorenzo Caputi, C. Robin Buell, Sarah E. O’Connor

## Abstract

Iridoids are specialized monoterpenes ancestral to asterid flowering plants (Albach *et al*, 2001; Stull *et al*, 2018). Iridoids play key roles in plant defense and are also essential precursors for pharmacologically important alkaloids (Dobler *et al*, 2011; Eisner, 1964). The biosynthesis of all iridoids involves the cyclization of a reactive enol intermediate. While this cyclization occurs spontaneously at low yields, it has long been hypothesized that a dedicated enzyme is involved in this process (Geu-Flores *et al*, 2012; Lichman *et al*, 2019b). Here, we report the discovery of asterid iridoid cyclases (ICYC). We show that these enzymes catalyze cyclization of the reactive intermediate to form the two major iridoid stereoisomers found in plants. Our work uncovers the last missing key step in the otherwise well-characterized iridoid biosynthesis pathway in asterids. This discovery unlocks the possibility to generate previously inaccessible iridoid stereoisomers, which will enable metabolic engineering for the sustainable production of valuable iridoid and iridoid-derived compounds.

## Main

Iridoids are widespread bicyclic monoterpenes found primarily in asterid plants (Albach *et al*., 2001; Burse & Boland, 2017; Kollner *et al*, 2022; Morita *et al*, 2003; Stull *et al*., 2018). Iridoids play important roles in plant defense; volatile iridoids are used by plants to repel or attract insects, whereas glycosylated forms serve as feeding deterrents (Birkett *et al*, 2011; Dinda *et al*, 2007; Dobler *et al*., 2011; Eisner, 1964; Koudounas *et al*, 2015). From a pharmacological perspective, iridoids possess promising anti-inflammatory activity (Viljoen *et al*, 2012), and moreover, serve as precursors for medicinally important monoterpenoid indole and ipecac alkaloids, which include anti-cancer (camptothecin and vinblastine), anti-malarial (quinine), putative anti-addiction (ibogaine), and emetic (emetine) agents (recently reviewed by (Lichman, 2021)).

Nepetalactol, which is the simplest iridoid and the common intermediate for all ca. 1000 known iridoids, is biosynthesized from geraniol-diphosphate (GPP). GPP is subjected to dephosphorylation by geraniol synthase (GES), hydroxylation by geraniol 8-hydroxylase (G8H) and oxidation by 8-hydroxygeraniol oxidase (8HGO) to yield 8-oxo-geranial (Collu *et al*, 2001; Miettinen *et al*, 2014; Simkin *et al*, 2013) (Fig. 1 a; Supplementary Fig. 1). The short chain dehydrogenase iridoid synthase (ISY) catalyzes a 1,4 reduction of 8-oxo-geranial to form 8-oxocitronellyl enol, which then cyclizes to form 7*S*, 4a*S*, 7a*R*-nepetalactol, along with a number of side products (Geu-Flores *et al*., 2012; Kries *et al*, 2017; Lichman *et al*, 2019a). Biosynthetic steps downstream of nepetalactol have been characterized in several plants, including *Catharanthus roseus* and *Camptotheca acuminata* (Supplementary Fig. 1) (Awadasseid *et al*, 2020; Irmler *et al*, 2008; Miettinen *et al*., 2014; Miller & Schuler, 2022; Murata *et al*, 2008).

**Fig. 1.**
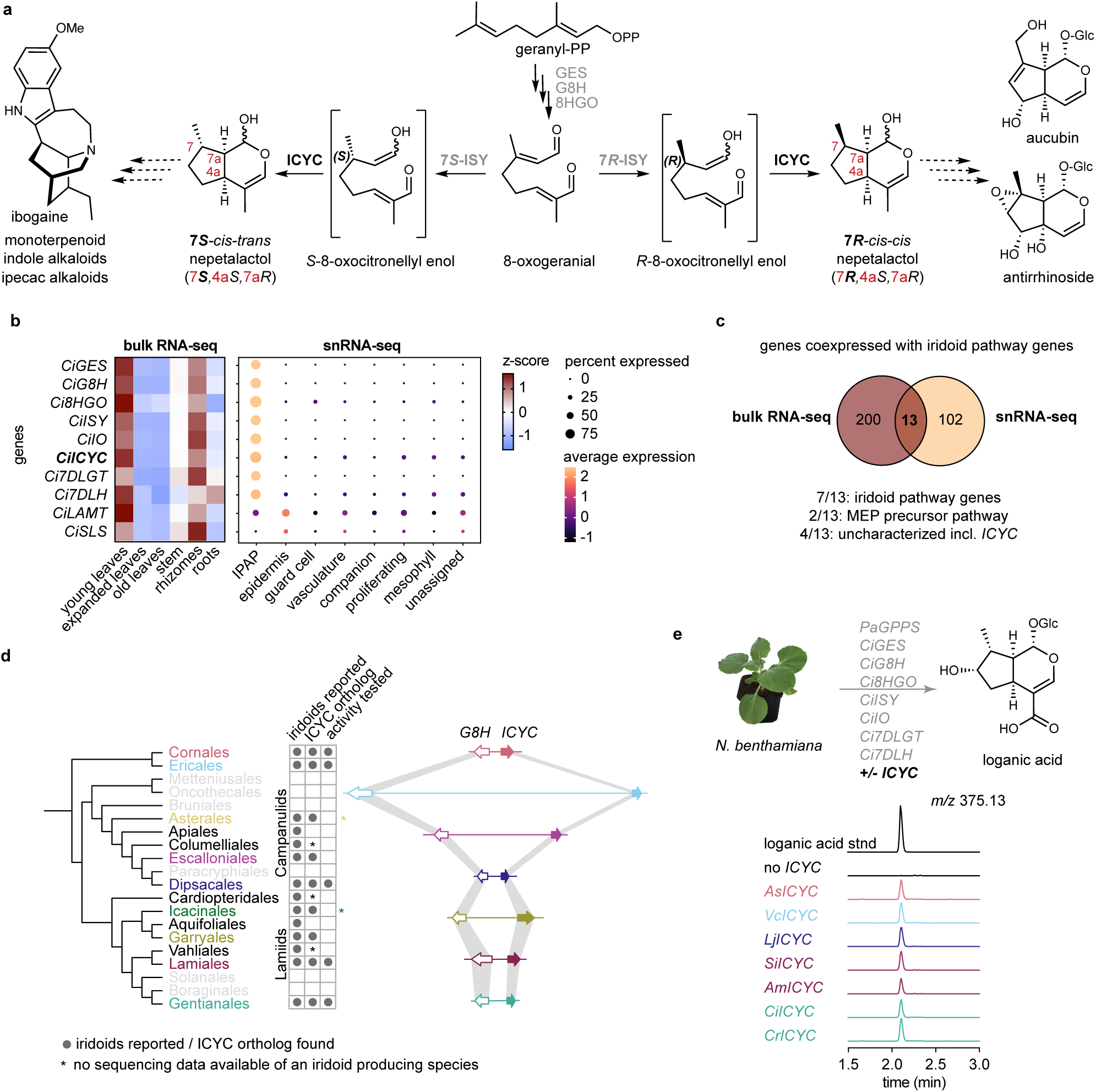
Identification of Iridoid cyclase (ICYC). **a**, Scheme showing iridoid biosynthesis including previously characterized iridoid pathway genes (gray). The 8-oxogeranial intermediate can be converted to either 7*S*-*cis-trans* nepetalactol or 7*R*-*cis-cis* nepetalactol by previously characterized species-specific stereoselective iridoid synthases (ISY) together with the newly identified iridoid cyclases (ICYC). For a complete scheme of the secoiridoid pathway including all intermediates see Supplementary Fig. 1. **b**, *Carapichea ipecacuanha* bulk tissue RNA-seq and single nuclei RNA-seq (snRNA-seq) show tight co-expression of orthologs of previously characterized secoiridoid pathway genes from *C. roseus* (Supplementary Fig. 3a) and the newly identified *CiICYC*. Complete tissue specific data is shown in Supplementary Fig. 3b. Cell clusters were grouped into cell types that were determined using marker genes (see Supplementary Fig. 4 for complete analysis of snRNA-seq dataset. **c**, Venn diagram showing overlay of co-expressed genes from both datasets. See Supplementary Fig. 5a for a detailed list of genes.) **d**, Identification of iridoid cyclase orthologs in various iridoid producing orders. The tree is based on the latest angiosperm phylogeny (Zuntini *et al*., 2024). Gray circles depict reported presence of iridoids within at least one species of the respective order. An asterisk depicts that no sequencing data was publicly available from a reported iridoid producing species from the respective order and thus ICYC presence could not be determined. On the right, a scheme depicting the synteny of the *ICYC* and *G8H* gene cluster. The synteny is shown for representative species of each order: *Alangium salviifolium* (Cornales), *Vaccinium corymbosum* (Ericales), *Escallonia rubra* (Escalloniales), *Lonicera japonica* (Dipsacales), *Eucommia ulmoides* (Garryales), *Antirrhinum majus* (Lamiales), *Carapichea ipecacuanha* (Gentianales) (Chanderbali *et al*, 2024; Li *et al*, 2019; Li *et al*, 2020; Yocca *et al*, 2023; Yu *et al*, 2022). The asterisk indicates that no genome data was available from an iridoid producing species of the Asterales and the Icacinales order. **e**, Representative *ICYC* orthologs from different orders all reconstituted the iridoid pathway up to loganic acid in *N. benthamiana*. Ci, *Carapichea ipecacuanha*; GES, geraniol synthase; G8H, geraniol hydroxylase; 8HGO, 8-hydroxygeraniol oxidase; ISY, iridoid synthase; IO, iridoid oxidase; ICYC, iridoid cyclase, 7DLGT, 7-deoxyloganetic acid glycosyl transferase; 7DLH, 7-deoxyloganic acid hydroxylase; LAMT, loganic acid methyltransferase; SLS, secologanin synthase; IPAP, internal phloem associated parenchyma; *As*, *Alangium salviifolium*; *Vc*, *Vaccinium corymbosum*; Lj, *Lonicera japonica*; Si, *Sesamum indicum*; Am, *Anthirrinium majus*; Ci, *Carapichea ipecacuanha*; Cr, *Catharanthus roseus*.

Of all possible nepetalactol stereoisomers, 7*S*-*cis-trans* (7*S*, 4a*S*, 7a*R*) and 7*R-cis-cis* (7*R*, 4a*S*, 7a*R*) nepetalactol are most commonly observed in nature (Fig. 1a) (Albach *et al*., 2001). 7*S*-*cis-trans* nepetalactol derived (“Route I”) iridoids, which are also precursors for alkaloid biosynthesis, are found among diverse asterid orders, whereas 7*R*-*cis-cis* nepetalactol derived (“Route II”) iridoids primarily occur in many Lamiales families (Albach *et al*., 2001; Jensen, 1991; Jensen, 1992). The stereoconfiguration at the C-7 position is set by a lineage-specific stereoselective ISY that generates either *S*-or *R*-8-oxocitronellyl enol (catalyzed by 7*S*-ISY and 7*R*-ISY, respectively) (Geu-Flores *et al*., 2012; Kries *et al*., 2017). Although 7*S*-*cis-trans* nepetalactol is observed in low yields when 7*S*-ISY is incubated with 8-oxogeranial, it has long been hypothesized that an additional enzyme assists the cyclization of 3*S*- or 3*R*-8-oxocitronellyl enol to 7*S*-*cis-trans* and 7*R-cis-cis* nepetalactol for the following reasons: (a) spontaneous cyclization occurs only *in vitro* and almost exclusively yields the *cis-trans* configuration, thus failing to account for the existence of the 7*R*-*cis-cis* configuration when 7*R*-ISY is used with 8-oxogeranial (Kries *et al*., 2017); (b) the known biosynthetic enzymes (GES, G8H, 8HGO, ISY) are not sufficient to reconstitute nepetalactol biosynthesis in *N. benthamiana* (Miettinen *et al*., 2014); and (c) recent work on *Nepeta*, a *Lamiaceae* genus that independently evolved iridoid biosynthesis, identified *Nepeta*-specific proteins that assist formation of the 7*S*-*cis-trans* nepetalactol scaffold from *S*-8-oxocitronellyl enol (Hernandez Lozada *et al*, 2022; Lichman *et al*, 2020; Lichman *et al*., 2019a). Indeed, the inclusion of a *Nepeta* Major Latex Protein Like (MLPL) enabled the successful pathway reconstitution of 7*S*-iridoids and downstream alkaloids in yeast and *N. benthamiana* (Dudley *et al*, 2022; Zhang *et al*, 2022).

Identifying the asterid iridoid cyclase(s) responsible for formation of 7*S*-*cis-trans* and 7*R-cis-cis* nepetalactol has been a long-standing challenge. The cyclization of this enol species to form the bicyclic scaffold that characterizes nepetalactol is an unusual chemical transformation that could be catalyzed by any number of protein scaffolds. To obtain a refined pool of gene candidates that might be responsible for this biosynthetic step, we generated *de novo* genome assemblies and high-resolution expression data, including single cell transcriptomics, of two evolutionarily distant members of the asterid clade, *Alangium salviifolium* (Cornales) and *Carapichea ipecacuanha* (Gentianales) (Supplementary Fig. 2; Supplementary tables 1-5) (Colinas et al. 2025). We identified highly conserved orthologs of *C. roseus* secoiridoid pathway genes in both species, confirming that iridoid biosynthesis is ancestral to asterids (Supplementary Fig. 3a). We next performed co-expression analysis using tissue-specific RNA-seq data, and found iridoid pathway genes to be tightly co-expressed in *C. ipecacuanha* young leaves and rhizomes and in *A. salviifolium* roots (Fig. 1b; Supplementary Fig. 3b, c) (Colinas et al. 2025). Additionally, we obtained single nuclei RNA-seq (snRNA-seq) data of *C. ipecacuanha* young leaves and constructed gene co-expression networks on the cell clusters resulting from the analysis of these data (Fig. 1b; Supplementary Fig. 4, 5; Supplementary Dataset 1). Iridoid pathway orthologs up to *7-DEOXYLOGANIC ACID HYDROXYLASE* (*7DLH*) are tightly co-expressed in a cell cluster corresponding to the Internal Phloem Associated Parenchyma (IPAP) cells, a cell type that has been previously shown to harbor early and intermediate iridoid biosynthesis steps in *C. roseus* (Burlat *et al*, 2004); later iridoid pathway gene orthologs are expressed in cell clusters containing epidermis marker genes (Fig. 1b; Supplementary Fig. 5b). The observed compartmentalization of IPAP-specific iridoid biosynthesis and epidermis-specific downstream biosynthesis is shown here for the first time for a *Rubiaceae* species and is identical to that found in *C. roseus* (*Apocynaceae*), suggesting that this spatial organization is conserved between these iridoid-producing families (Burlat *et al*., 2004; Li *et al*, 2025; Li *et al*, 2023b; Miettinen *et al*., 2014; Sun *et al*, 2023).

With these datasets in hand, we filtered the transcript lists generated from bulk tissue RNA-seq and snRNA-seq co-expression analyses for high absolute expression levels (CPM > 50 in young leaves; cluster average expression > 1 within the IPAP cell cluster), based on the assumption that the cyclase gene would be highly expressed. Combining the filtered lists provided 13 gene candidates (Fig. 1c; Supplementary Fig. 6a). These 13 genes included all seven IPAP-specific iridoid biosynthesis genes, two genes of the 2-*C*-methyl-D-erythritol 4-phosphate (MEP) pathway that makes the GPP precursor isopentenyl phosphate (IPP), and only four uncharacterized genes (Fig. 1c, Supplementary Fig. 6a). Interestingly, when less stringent absolute expression value cutoffs were applied, the overlaid list also contained orthologs of the three known *C. roseus* basic helix-loop-helix (bHLH) iridoid biosynthesis (BIS) transcriptional regulators, highlighting that this strategy can also be used to identify cell type specific transcriptional regulators (Supplementary Fig. 6b) (Colinas *et al*, 2021; Van Moerkercke *et al*, 2016; Van Moerkercke *et al*, 2015).

We had previously developed a mass spectrometry-based detection method for 7*S*-*cis-trans* nepetalactol derived iridoids in transfected *N. benthamiana* leaves (Dudley *et al*., 2022). We capitalized on this method to screen the activity of the cyclase gene candidates by assaying them in the context of upstream (*GPPS*, *GES*, *G8H*, *8HGO*, *ISY*) and downstream (*7-DEOXYLOGANETIC ACID GLUCOSYL TRANSFERASE* (*7DLGT*), *7DLH*)) iridoid biosynthetic pathway genes. The cyclase candidates were expressed with *C. ipecacuanha* orthologs of iridoid biosynthetic genes predicted to generate loganic acid, an iridoid derived from 7*S*-*cis-trans* nepetalactol, in *N. benthamiana*. Inclusion of one of the candidates (Lemenager *et al*, 2005), which was functionally annotated as a methyl esterase, resulted in the efficient production of loganic acid (Fig. 1e). The enzyme was thus named Iridoid cyclase (ICYC). This protein is entirely unrelated to the *Nepeta*-specific cyclases, clearly indicating that iridoid cyclases arose convergently at least once. We systematically compared the CiICYC sequence against available data for 20 asterid clades (Zuntini *et al*, 2024) and identified ICYC orthologs in all clades reported to produce iridoids with the exception of the Apiales and Aquifoliales iridoid producing genera *Griselinia* and *Helwingia*, respectively (Jensen & Nielsen, 1980; Murayama *et al*, 2004). Analogously, an ICYC ortholog appeared to be absent in non-iridoid producing clades (Fig. 1d; see Supplementary Fig.7 for a tree with all orthologs). Interestingly, *ICYC* is located next to the iridoid pathway gene *G8H*, forming a small biosynthetic gene cluster that is conserved in all iridoid producing orders for which genome data is available (Fig. 1d).

In addition to *C. ipecacuanha* ICYC (CiIYC, lamiids, Gentianales, Rubiaceae), we selected orthologs from six additional species representing different asterid orders and families: the 7*S-cis-trans* nepetalactol producing species *A. salviifolium* (AsICYC, Cornales), *Vaccinium corymbosum* blueberry (VcICYC, Ericales), *Lonicera japonica* (LjICYC, campanulids, Dipsacales), and *C. roseus* (CrICYC, lamiids, Gentianales, Aponcynaceae), and the 7*R-cis-cis* nepetalactol producing Lamiales species *Sesamum indicum* (SiICYC, Pedaliaceae) and *Antirrhinum majus* (AmICYC, Plantaginaceae) (Colinas *et al*, 2025; Fuji *et al*, 2018; Kakuda *et al*, 2000; Kries *et al*., 2017; Lawas *et al*, 2023). Inclusion of each of these genes resulted in the effective reconstitution of loganic acid biosynthesis in *N. benthamiana* (Fig. 1e, Supplementary Fig. 8). We further used these cyclases to reconstitute the complete downstream *C. ipecacuanha* and *A. salviifolium* iridoid pathways (Supplementary Fig. 9). Finally, to confirm ICYC activity in a native plant, we performed virus induced gene silencing (VIGS) of *CrICYC* in *C. roseus* (a technique not available in *C. ipecacuanaha*) and detected reduced iridoid (secologanin) content, consistent with the proposed iridoid cyclization function (Supplementary Fig. 10).

To rigorously characterize the direct product of ICYC, we assayed seven recombinantly produced ICYC orthologs together with 8-oxogeranial and the previously characterized 7*S*-ISY from *C. roseus* (CrISY) and 7*R*-ISY from *A. maju*s (AmISY) (Fig. 2; Supplementary Fig. 11-12) (Geu-Flores *et al*., 2012; Kries *et al*., 2017). CrISY and AmISY reduce 8-oxogeranial to form the unstable intermediate *S*- or *R*-8-oxocitronellyl enol, respectively, which, in the absence of a cyclase, spontaneously forms a small amount of *cis-trans* nepetalactol, along with a wide variety of side products (Fig. 2a; see Supplementary Fig. 11 for detailed description of all spontaneous products) (Kries *et al*., 2017; Lichman *et al*., 2019a). When CrISY was assayed together with any of the ICYC orthologs, 7*S*-*cis-trans* (*7S*, 4a*S*, 7a*S*) nepetalactol accumulation was substantially increased compared to when CrISY was assayed alone (Fig. 2b). When AmISY was assayed with ICYC orthologs, a new peak appeared (Fig. 2c). We confirmed the identity of this ICYC specific product as 7*R-cis-cis* (*7S*, 4a*S*, 7a*S*) nepetalactol by chemical oxidation and comparison with an authentic standard of 7*R-cis-cis* nepetalactone (Fig. 2d-f) (Hernandez Lozada *et al*., 2022). Additionally, we observed comparable results when we assayed the ICYC orthologs directly with *S*-8-oxocitronellal and *R*-8-oxocitronellal at high concentrations of general acid (0.5 M MOPS), conditions that promote partial tautomerization to the putative cyclase substrate, 8-oxocitronellyl enol (Lichman *et al*., 2019a) (Supplementary Fig. 13). Taken together, these results show that ICYCs exclusively produce 4a*S*, 7a*S* nepetalactol regardless of the stereochemistry of the C-7-position. Strikingly, while all ICYC orthologs generate *7S*-*cis-trans* nepetalactol from *S*-8-oxocitronellal, 7*R*-*cis-cis* nepetalactol is most efficiently generated when ICYC orthologs from the Lamiales species known to produce this stereoisomer (i.e. AmICYC and SiICYC) are used. This observation suggests that enzymes in these species might have specialized to adapt to the 7*R* substrate. We also assayed the product profile when ICYC was added after ISY reduced 8-oxogeranial to 8-oxocitronellyl enol (Supplementary Fig. 14a). Under these conditions, no increase in nepetalactol is observed, suggesting that 8-oxocitronellyl enol must be immediately transferred to ICYC after it is formed by ISY. In support of this hypothesis, we found that ISY and ICYC interact (as measured by split luciferase assays), suggesting that substrate channeling, which would protect the reactive enol species, could take place between these two proteins (Supplementary Fig. 14b).

**Fig. 2.**
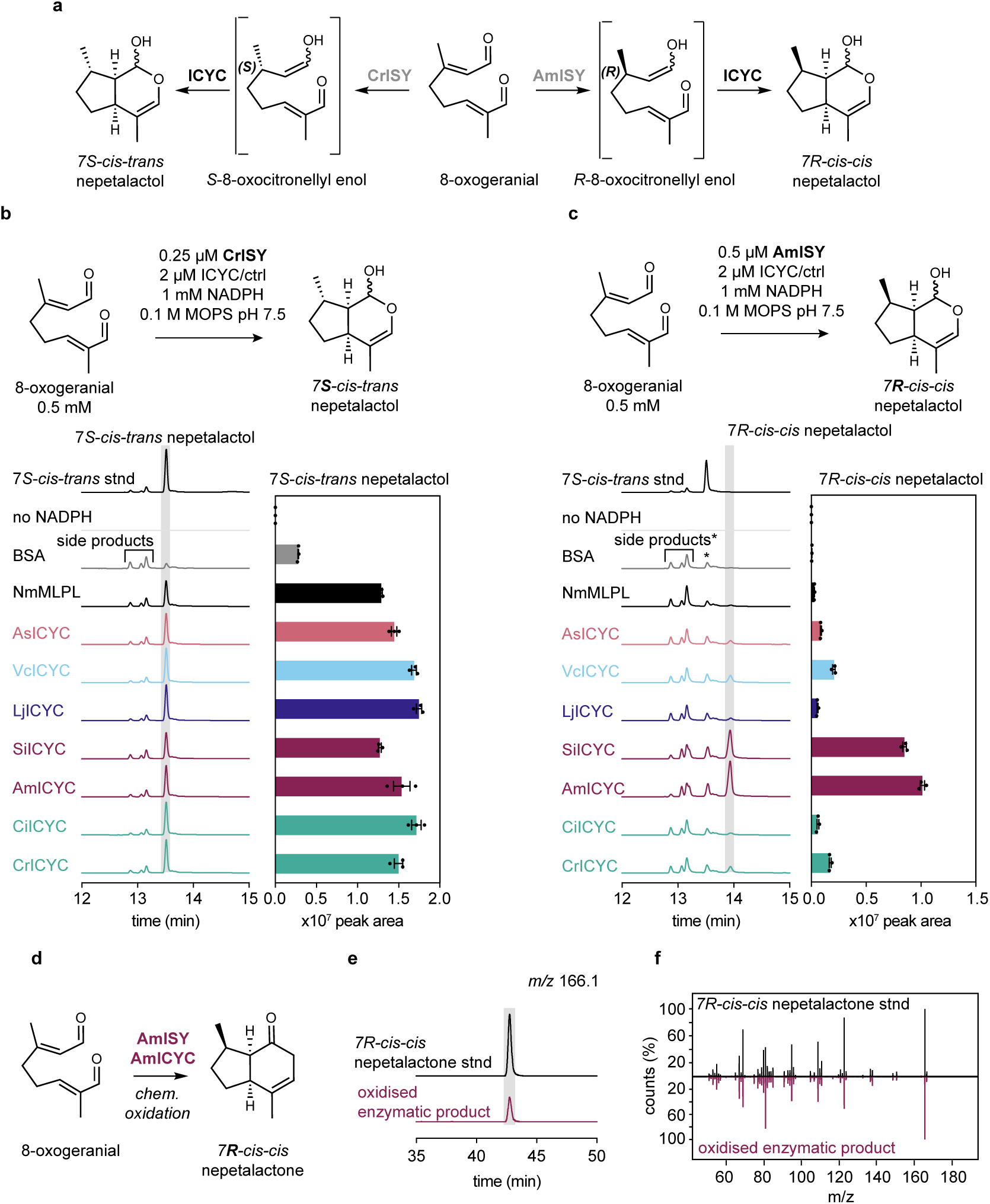
In vitro activity assays reveal stereoselectivity of ICYCs. **a**, Scheme depicting reaction products in of 7*S* and 7*R* stereo-selective ISY together with ICYC. **b**, **c** Assays were performed as indicated and analyzed by gas chromatography mass spectrometry (GC-MS). Bovine serum albumin (BSA) was used as a negative control. Displayed chromatograms are total ion chromatograms of one representative replicate. See Supplementary Fig. 11 for detailed descriptions of chromatograms including side products and Supplementary Fig. 12 for fragmentation patterns of standards and enzymatic products. Bar graphs depict peak areas from three replicates of nepetalactol (shaded in gray), error bars are standard error of the mean (SEM). **b**, Assays of ICYC orthologs with the 7*S* selective *C. roseus* ISY (CrISY) show a dramatic increase in *7S-cis-trans* nepetalactol in the presence of ICYC orthologs. **c**, Assays of ICYC orthologs with the 7*R* selective *A. majus* ISY (AmISY) reveal appearance of a peak consistent with 7*R*-*cis-cis* nepetalactol in presence of ICYC. The asterisk indicates that these side products are the 7*R* enantiomers that coelute with the 7*S* series when using an achiral stationary phase as used here (see Supplementary Fig. 11). **d**, To confirm the identity of the AmISY-AmICYC enzymatic product, five enzymatic reactions were pooled and chemically oxidized to 7*R*-*cis-cis* nepetalactone (see methods). **e**, Chiral GC-MS analysis of the chemically oxidized product showed that it had had the same retention time and the identical mass (extracted ion chromatogram for nepetalactone mass *m/z* 166.1) as the authentic standard (Hernandez Lozada *et al*., 2022) on a chiral GC column. **f**, The MS fragmentation pattern of the oxidized enzymatic product was identical to the standard confirming that the enzymatic product is indeed 7*R*-*cis-cis* nepetalactol.

ICYC is a methyl esterase (MES) type *α*/*β* hydrolase. These methyl esterases are ubiquitously found in plants, but to the best of our knowledge have never previously been associated with non-esterase functions such as cyclization. Many methylesterase family members play roles in activating the plant hormones methyl jasmonate, methyl salicylate or indole-3-acetic acid methyl ester *via* demethylation whereas other members act as esterases in monoterpene indole or ipecac alkaloid biosynthetic pathways (Fig. 3a; Supplementary Fig. 15) (Colinas *et al*, 2024; Dogru *et al*, 2000; Farrow *et al*, 2019; Forouhar *et al*, 2005; Hong *et al*, 2022; Trenti *et al*, 2021; Vlot *et al*, 2008; Volk *et al*, 2019; Yang *et al*, 2008). All *α*/*β* hydrolases utilize a conserved catalytic triad (serine, aspartate/glutamate and histidine) along with a glycine-rich oxyanion hole to catalyze substrate hydrolysis (Fig. 3a, b) (Ollis *et al*, 1992; Rauwerdink & Kazlauskas, 2015). Although ICYC orthologs form a phylogenetic clade well separated from all other methylesterases, indicating a monophyletic origin (Fig. 3a; Supplementary Fig. 15), the catalytic triad and oxyanion hole motifs are still present, as in canonical esterases (Fig. 3a; Supplementary Fig. 16). No obvious differences were noted within the active sites of cyclases from species that produce 7*R* nepetalactol compared to those that produce 7*S* nepetalactol (Supplementary Fig. 17). All ICYC orthologs showed esterase activity when tested with the model substrate 4-nitrophenyl acetate, indicating that these catalytic motifs remain functional (Fig. 3a; Supplementary Fig. 18a). To assess which residues are responsible for catalysis of cyclization, we docked the *S*-8-oxocitronellyl enol substrate to a CiICYC alphafold3 model with AutoDock Vina to pinpoint residues in the binding pocket. We also used our phylogenetic analysis to select amino acid residues that are conserved in ICYC orthologs (Fig. 3a,b). Unsurprisingly, mutations of any of the catalytic triad amino acid in CiICYC (S83A; D210A; H238A) abolished esterase activity (Supplementary Fig. 18b) (Mattern-Dogru *et al*, 2002). D210A was largely insoluble and could therefore not be assayed reliably. When these mutants were tested for cyclase activity *in vitro* (Fig. 3c) and in *N. benthamiana* (Fig. 3d), D210A and H238A mutants were inactive, whereas the S83A mutant exhibited some cyclization activity. Out of the four additional mutants within the putative cyclase active site, V15G and W133A led to reduced amounts of the nepetalactol product (Fig. 3c, d). Although more extensive experimentation is required to understand the mechanism of this unusual cyclization (Supplementary Fig. 19a-c), we speculate that these residues shape the binding pocket of ICYC, allowing 8-oxocitronellyl enol to adopt the conformation required for cyclization. The amino acid H238, mutation of which abolished detectable activity, may play a role in orienting the substrate or, alternatively, it could interact with the enol moiety of 8-oxocitronellyl enol, which could in turn activate the substrate to undergo stepwise cyclization (Supplementary Fig. 19d) (Lindner *et al*, 2014).

**Fig. 3.**
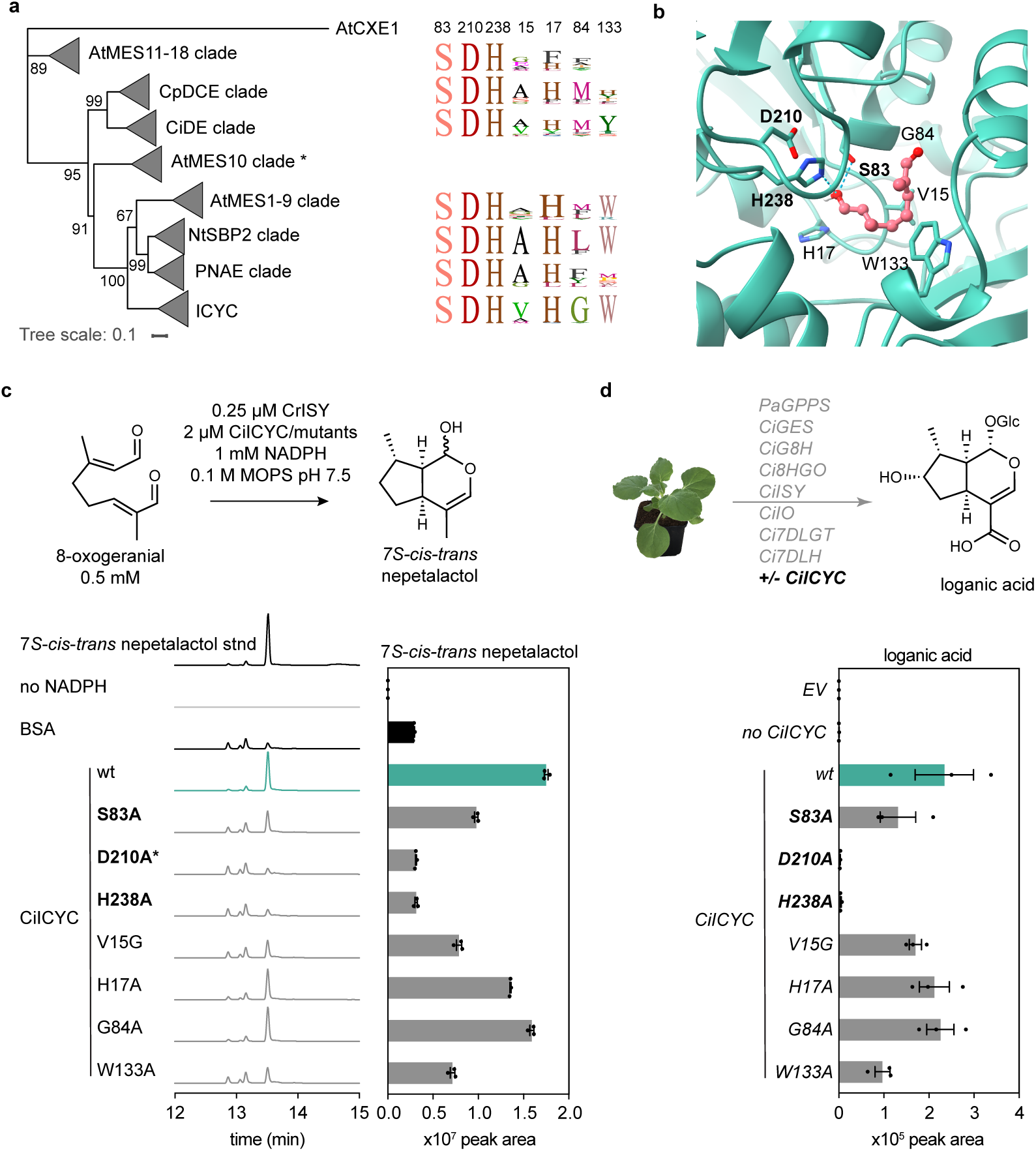
ICYC is a member of the methylesterase family. **a**, A condensed phylogenetic tree showing that ICYC is related to methylesterases that esterify substrates in plant hormone activation pathways and in alkaloid biosynthesis. Each clade is named after a member found within the respective clade. See Supplementary Fig. 16 for an extended version of this tree. Sequence data highlights the conservation of the catalytic triad and other amino acid residues that were subjected to mutation in CiICYC. The asterisk indicates that AtMES10 was a single member clade, thus no sequence logos were generated. At, *Arabidopsis thaliana*; CXE, Carboxylesterases; CiDE, *Carapichea ipecacuanha* deacetyl(iso)ipecoside esterase; CpDCE, *Cinchona pubescens* dihydrocorynantheine aldehyde esterase; RsPNAE, *Rauvolfia serpentina* polyneuridine aldehyde esterase. **b**, Putative active site of an alphafold3 structural model of CiICYC with the docked (AutoDock Vina) substrate *S*-8-oxocitronellyl enol substrate. Indicated amino acids were chosen for mutagenesis studies because they are part of the catalytic triad (in bold) or are in proximity to the docked substrate and are conserved in ICYC orthologs. **c**, *in vitro* assays show that D210A and H238A ICYC mutants yield a reaction profile that is identical to the negative control. S83A, V15G and W133A mutants show reduced levels of nepetalactol compared to wild-type ICYC, whereas H17A and G84A mutants behaved similarly to wild-type (wt) ICYC. The asterisk indicates that the D210A mutant exhibited poor solubility and thus the assays with this mutant result should be interpreted cautiously. **d**, Reconstitution of loganic acid biosynthesis with CiICYC and CiICYC mutants yielded results that are consistent with results from *in vitro* assays shown in **d**. The discovery of these stereoselective ICYC orthologs fully explains the dominance of 7*S cis-tran*s and 7*R cis-cis* nepetalactol derived iridoids in the asterids. Although further studies are required to fully reveal the mechanistic basis of this cyclization (Lichman *et al*., 2019b; Lindner *et al*., 2014), this work clearly shows how a widely present methyl esterase has been coopted throughout the asterids– a clade comprising over 100 plant families– to perform this unusual reaction. This discovery unlocks the possibility for formation of previously inaccessible iridoid stereoisomers, which will enable metabolic engineering for sustainable production valuable iridoid and iridoid derived compounds.

## Acknowledgements

We thank the gardeners Eva Rothe and Elke Goschala for growing and maintaining *A. salviifolium* and *C. ipecacuanha* plants as well as Franz Kaltofen for growing *N. benthamiana* plants. We thank Benjamin Lichman for providing pOPINF_NmMLPL construct, Quentin M. Dudley for providing *NmMLPL* and *GPPS* constructs, and Prashant Sonawane for providing the modified 3Ω1 vector. We are grateful to Néstor J. Hernández Lozada for providing *7R-cis-cis* nepetalactone standard and Carlos E. Rodríguez-López for providing *S*-8-oxocitronellal and *R*-8-oxocitronellal. We thank Helena Leucke for assistance with RNA extraction, Maritta Kunert for assistance with UPLC-MS/MS and GC-MS analyses and Yoko Nakamura for NMR structural confirmation of commercial 7*S*-*cis*-*trans* nepetalactol standard. We thank Lemor Carlton for assistance with nanopore cDNA sequencing. We thank Brieanne Vailllancourt for assistance with sequencing data management. We are grateful to the Max Planck Society for funding and Horizon 2020 (MIAMi, grant no. 814645) and the Leibniz Prize, Deutsche Forschungsgemeinschaft (DFG, German Research Foundation) – 505457618 awarded to Sarah E. O’Connor. We acknowledge funding from the University of Georgia, Georgia Research Alliance, and Georgia Seed Development to C. Robin Buell.

## Author contributions

M.C. designed all experimental work and analyzed data. R.D. performed RNA extraction, Illumina sequencing genome sequencing and initial genome assemblies. J.P.H. performed genome annotation. M.C. processed tissue-specific gene expression data and performed co-expression analyses. J.C.W. performed snRNA-seq and data processing. K.G. performed VIGS and qPCRs assisted by L.C. R.M.A. performed chemical oxidation of 7-*R-cis-cis* nepetalactol enzymatic product. C.M. performed split-Luciferase assays. C.T., J.W. and A.D. performed cloning. J.W. and A.D. performed recombinant protein purifications. M.C., C.T., and J.W. performed reconstitution in *N. benthamiana*. M.C. performed *in vitro* enzyme assays. S.H. developed the UPLC-MS/MS method. G.R.T. developed the GC-MS method. A.A.L provided *C. ipecacuanha* living specimens. M.C., C.R.B. and S.E.O. designed the study. M.C. and S.E.O. wrote the manuscript with input from all other authors.

## Data availability statement

All sequencing data associated with this study are available at the National Center for Biotechnology Institute Sequence Read Archive BioProject PRJNA1270996 (to be made public upon acceptance).

## Methods

### Plant sampling and RNA extraction

A. C. ipecacuanha and A. salviifolium plants were previously obtained as described (Colinas et al. 2025). Plants labelled with different numbers refer to independent individual plants. RNA-seq data from plants labelled as “plant 1” was previously published (NCBI BioProject PRJNA1169657). All plants were grown under the following conditions: 12/12 h light/dark 28-30°C/24-26°C, 70-80% humidity. Plants for Illumina sequencing and expression analyses (see Supplementary Fig. 2 a, b for photographs): C. ipecacuanha plants labelled as “plant 2” and “plant 3” were both 1.5 years old; A. salviifolium plants labelled as “plant 2” and “plant 3” were 2.5, or 4 years old, respectively. Dissected tissues from C. ipecacuanha “plant 2 and 3” and A. salviifolium “plant 2” were flash frozen and shipped on dry ice to FutureGenomics for RNA extraction and sequencing (see below). Additional tissues were harvested for Oxford Nanopore Technologies (ONT) sequencing for genome annotation (young leaves, mature leaves, green stem and roots of 2-year-old C. ipecacuanha plants; leaf buds, young leaves, mature leaves, roots of 3-year-old A. salviifolium plants). RNA of samples for ONT sequencing and A. salviifolium “plant 3” samples was extracted in-house as follows. Dissected tissues were immediately flash frozen into liquid nitrogen and ground in liquid nitrogen using an IKA A11 basic analytical mill or mortar and pestle. Total RNA was extracted using the RNeasy Plant Mini Kit (Qiagen) according to the manufacturer’s instructions, including on column DNAse digest. RNA concentrations and purity were determined with a Nanophotometer N60 (Implen).

### RNA Illumina sequencing

Illumina sequencing of *C. ipecacuanha* “plant 2 and 3” and *A. salviifolium* “plant 2” was performed at FutureGenomics as follows. Flash-frozen plant tissues were powdered in liquid nitrogen using mortar and pestle and RNA was extracted using Zymo Quick-RNA Plant Kit (Zymo Research, CA, USA). The quality of the RNA was analyzed using RNA ScreenTape on an Agilent 4200 TapeStation System (Agilent Technologies Netherlands BV, Amstelveen, The Netherlands) and the quantity was measured using a Qubit 3.0 Fluorometer (Life Technologies Europe BV, Bleiswijk, The Netherlands). Illumina RNAseq libraries were prepared using NEBNext Ultra II Directional RNA Library Prep Kit Illumina and NEBNext Multiplex Oligos for Illumina (New England Biolabs, MA, USA). The quality of the libraries was checked using Agilent D1000 screentape on an Agilent 4200 TapeStation System. RNA library paired-end sequencing (2x150 bp) was performed using Illumina’s NovaSeq 6000 technology. Two rounds of sequencing were carried out to reach the desired output target.

Illumina sequencing of *A. salviifolium* “plant 3” was performed at Novogene. RNA samples were shipped to Novogene where mRNA library preparation and sequencing were performed according to the company’s standard protocol for mRNA sequencing. RNA integrity and quantitation were assessed on a Bioanalyzer (Agilent Technologies, CA, USA). All samples were above the required minimum RIN value. Sequencing was performed on an Illumina NovaSeq X Plus PE150 platform with a data output target of 9 G of raw data.

### Oxford Nanopore Technologies full-length cDNA sequencing

mRNA was purified from the total RNA using the Dynbeads mRNA Purification Kit (Invitrogen, MA, USA) and input into the Oxford Nanpore Technologies (ONT) SQK-PCS111 kit to generate fl-cDNA libraries. Resultant libraries were sequenced on a MIN106 Rev. D flowcell before being basecalled using Guppy (v6.5.7) (https://nanoporetech.com/software/other/guppy) using the super high accuracy model (dna_r9.4.1_450bps_sup.cfg) and the parameters -q 0, –trim_strategy none, and --calib-detect.

### Genome sequencing and assembly

Nuclei were extracted from young leaves using a nuclei isolation protocol (Carrier *et al*, 2011). The nuclear pellet was resuspended in Qiagen buffer G2 with RNaseA and proteinaseK, and DNA extraction was further continued according to the instructions of the Qiagen Genomic Tip/100G protocol (Qiagen, Venlo, The Netherlands). The quality of the DNA was analyzed using Genomic DNA ScreenTape on an Agilent 4200 TapeStation System (Agilent Technologies Netherlands BV, Amstelveen, The Netherlands) and the quantity was measured using a Qubit 3.0 Fluorometer (Life Technologies Europe BV, Bleiswijk, The Netherlands). Nanopore sequencing libraries were prepared using the Ligation Sequencing Kit V14 (SQK-LSK114) according to the manufacturer’s instructions (Oxford Nanopore Technologies, Oxford, UK). Each library was run on an R10.4.1 PromethION flowcell (FLO-PRO114M; Oxford Nanopore Technologies (ONT), Oxford, UK) and reloaded on a daily basis after a nuclease flush with Flow Cell wash kit (EXP-WSH004). Super-accuracy basecalling was done using Guppy 6.3.2. Illumina DNAseq libraries were prepared using the Nextera Flex kit according to the manufacturer’s instructions (Illumina, San Diego, CA, USA) and were sequenced in paired-end mode (2 × 150 bp) using Illumina’s NovaSeq 6000 technology.

ONT genomic reads were assembled using Flye (v.2.9.1), with settings “overlap 10K, error rate 0.025, no-alt-contigs” (Kolmogorov *et al*, 2019). The genome contigs were polished using nanopore reads by Medaka (https://github.com/nanoporetech/medaka) and then polished twice using Illumina reads by Pilon (v.1.23) (Walker *et al*, 2014). The polished genome sequence was then collapsed using Purge dups (Guan *et al*, 2020).

### Genome annotation

The genome assemblies were repeat masked by first creating a custom repeat library (CRL) for each genome. Repeats were first identified with RepeatModeler (Flynn *et al*, 2020) (v2.03) and protein coding genes filtered out from the repeat database using ProtExcluder (Campbell *et al*, 2014) (v1.2) to create a CRL. The CRL was then combined with Viridiplantae repeats from RepBase (v20150807) to generate the final CRL for each genome. Each genome assembly was repeat-masked using the respective final CRL and RepeatMasker (Chen, 2004) (v4.1.2-p1) using the parameters -e ncbi -s - nolow -no_is -gff.

RNA-seq libraries were processed for genome annotation by first cleaning with Cutadapt (Martin, 2011) (v2.10) using a minimum length of 100 nt and quality cutoff of 10 then aligning the cleaned reads to the respective genome using HISAT2 (Kim *et al*, 2019) (2.1.0). Oxford Nanopore (ONT) cDNA reads were processed with Pychopper (https://github.com/epi2me-labs/pychopper) (v2.7.10) and trimmed reads greater than 500 nt were aligned to the respective genome using minimap2 (Li, 2021) (v2.17-r941) with a maximum intron length of 5,000 nt. The aligned RNA-seq and ONT cDNA reads were each assembled using Stringtie (Kovaka *et al*, 2019) (v2.2.1) and transcripts less than 500 nt were removed.

The initial gene models for each genome were created using BRAKER2 (Brůna *et al*, 2021) (v2.1.6) using the soft-masked genome assemblies and the aligned RNA-seq libraries as hints. The gene models were then refined using two rounds of PASA2 (Haas *et al*, 2003) (v2.5.2) to create a working gene model set for each genome. High-confidence gene models were identified from each working gene model by filtering out gene models without expression evidence, or a PFAM domain match, or were a partial gene model or contained an interior stop codon. Functional annotation was assigned to by the searching the working gene models proteins against the TAIR (Lamesch *et al*, 2012) (v10) database and the Swiss-Prot plant proteins (release 2015_08) database using BLASTP (Altschul *et al*, 1990) (v2.12.0) and the PFAM (Mistry *et al*, 2021) (v35.0) database using PfamScan (Li *et al*, 2015) (v1.6) and assigning the annotation based on the first significant hit.

### Single nuclei RNA-seq of *C. ipecacuanha* young leaves

Nuclei were isolated following the protocol outlined by (Li *et al*, 2022). For *C. ipecacuahna* young leaves 0.01% Triton X-100 was used and RNase inhibitor (Sigma Protector RNase Inhibitor, Cat. No. 3335402001; SigmaAldrich, MO, USA) was added to the nuclei isolation buffer for a final concentration of 0.5U/μl. Nuclei were stained with DAPI (4’,6-diamidino-2-phenylindole) and sorted using Fluorescent Activated Cell Sorting (FACs). Nuclei were concentrated by spinning at 300G for 5 min. Concentrated nuclei were used for single cell RNAseq library construction using the PIPseq T20 v4.0Plus Kit (Illumina, CA, USA) with 1μl of additional RNase inhibitor.

### Single cell transcriptomics and co-expression analyses

Single nuclei RNA-seq reads were processed using the “barcode” command from pipseeker-v3.1.3 (Illumina, CA, USA). The pipseeker processed fastq files and the generated barcode whitelist were used as input into the STARsolo (v2.7.10b) (Kaminow *et al*, 2021) alignment program. The following parameters were employed; --alignIntronMax 5000, --soloUMIlen 12, --soloCellFilter EmptyyDrops_CR, --soloFeatures GeneFull, --soloMultiMappers EM, --soloType CB_UMI_Simple. Seurat v4.3.0.1 was used for downstream analysis. Samples were filtered to retain high quality cells by removing cells with less than 300 genes or more than 10,000 genes and less than 500 UMI’s or more than 30,000 UMI’s. Samples were also run through DoubletFinder (McGinnis *et al*, 2019) to remove suspected doublets. Reciprocal PCA (RPCA) was used to integrate the two replicates together using the top 3,000 variable genes. The top 60 principal components were used with a resolution parameter of 0.5 to calculate the Uniform manifold approximation and projection (UMAP).

SimpleTidy_GeneCoEx (Li *et al*, 2023a) was used to for co-expression analysis. An R value cutoff of 0.7 with a resolution parameter of 3 was used to generate 27 modules containing 5 or more genes, which comprised 29,765 genes total. Module 16 contained IPAP-specific genes of interest (1,496 genes in module).

### Mapping of bulk RNA-seq to *A. salviifolium* and *C. ipecacuanha* assembled genomes

Adapter-cleaved raw fastq files were received and reads were quality-checked with FastQC and trimmed using Trimmomatic (Bolger *et al*, 2014) on an in-house Galaxy server (Afgan *et al*, 2018). Reads belonging to the same sample but obtained in two sequencing rounds were concatenated. Reads were mapped to genomes using CLC Genomics workbench 21.0.4 (Qiagen) with these parameters: mismatch cost, 2; insertion cost, 3; deletion cost, 3; length fraction, 0.85; similarity fraction 0.9; auto-detect paired distances, on; maximum number of hits for a read, 20. Expression values are unique counts per gene. TMM normalized CPM values were used for downstream analyses. To identify iridoid pathway genes in *C. ipecacuanha* and *A. salviifolium* amino acid sequences of the characterized iridoid pathway enzymes from *C. roseus* were blasted (tblastn) against the transcript working models derived from the genomes. The highest blast hit for each gene was chosen for expression analysis and cloning (Supplementary Fig. 3a). Blasting of enzymes known to perform cyclizations in *Nepeta* (Lichman *et al*., 2020; Lichman *et al*., 2019a) yielded poorly conserved sequences (max. 59 % identity) that were not co-expressed with iridoid pathway genes, and were thus not considered as orthologs. *C. ipecacuanh*a tissue-specific co-expression analysis was performed on an expression atlas containing data from all tissues from “plants 1-3”. *CiISY* expression was used as a bait for Pearson correlation.

### Gene cloning

cDNA was prepared from total RNA of *A. salviifolium* leaf buds and roots, *C. ipecacuanha* young leaves and *C. roseus* leaves (extracted as described above) using the iscript cDNA Synthesis Kit (Biorad) according to manufacturer’s instructions. *NmMLPL* and *PaGPPS* genes had been cloned previously (Dudley *et al*., 2022; Lichman *et al*., 2020). Genes from all other species and *CiICYC* protein mutants were obtained as synthetic sequences from Twist Biosciences. *CDS* sequences were amplified with the Q5 High-Fidelity 2X Master Mix (New England Biolabs) using cDNA or synthetic fragments as templates and gene specific primers containing overhangs for In-Phusion cloning (Supplementary Table 7). Amplified sequences were gel-purified using the Zymoclean Gel DNA Recovery Kit (Zymo Research) and cloned using the 5x In-Fusion Snap Assembly Master Mix (TaKaRa Bio). For pathway reconstitution in *N. benthamiana*, coding regions were inserted into a modified 3Ω1 vector (contains *UBQ10* promoter and terminator from *Solanum lycopersicum* (Cardenas *et al*, 2019)) previously digested with BsaI-HF v2 (New England Biolabs, NEB). For expression in *Escherichia coli CDS’* were cloned into pOPINF (Berrow *et al*, 2007) previously digested with KpnI-HF and HindIII-HF (NEB). For split-luciferase assays CiICYC and CiISY were cloned into KpnI-HF and SalI-HF (NEB) digested pCAMBIA1300-NLuc, and KpnI-HF and PstI-HF (NEB) digested pCAMBIA1300-Cluc (Chen *et al*, 2008). Cloning reactions were transformed into heat shock competent *E. coli* TOP10 and grown over night in a 37°C incubator on LB agar plates containing the respective antibiotics. Plasmids were isolated from overnight cultures of single colonies using the Wizard Plus SV Minipreps DNA Purification System kit (Promega) and inserted sequences confirmed by Sanger sequencing.

### *A. tumefaciens* mediated transient expression in *N. benthamiana*

*A. tumefaciens* GV3101 cells were transformed through electroporation, recovered in YEB without antibiotics and incubated on YEB plates containing antibiotics (rifampicin and gentamycin and the appropriate antibiotic for plasmid selection) at 28°C for 48 hours. Colony PCR was done on single colonies to confirm presence of plasmids. Positive colonies were grown in liquid YEB for 24 hours. From these cultures glycerol stocks were prepared and stored at –80°C. Three-four week-old *N. benthamiana* plants (grown in a greenhouse with 16h/8h light/dark and 23-26°C /16-22°C, 40-70% humidity) were used for agroinfiltration as previously described (Colinas *et al*., 2025; Zhang *et al*, 2020): cells from glycerol stocks were spread on YEB plates containing antibiotics and 100 µM acetosyringone and grown for 24 hours until a visible layer of bacteria appeared. The bacteria were transferred to 1-2 ml of infiltration medium (10 mM MES, 10 mM MgCl_2_, 100 μM acetosyringone, pH 5.7), gently resuspended and the OD_600_ measured in 1:10 dilutions using an Implen OD600 DiluPhotometer. For pathway reconstitution experiments, strains were mixed and diluted in infiltration buffer to OD_600_=0.1 per strain. For split luciferase assays (see below), the strains were mixed at OD_600_=0.13 per strain. Strains harboring constructs with NLuc or CLuc fused to CiICYC of CiISY or empty vectors containing free NLuc or CLuc as controls were infiltrated in combinations as indicated. A strain harboring a construct with the *p19* gene was co-infiltrated in all cases. The culture mixtures were infiltrated into *N. benthamiana* leaves and grown for 5 days under grow lights (16h/8h light/dark). Replicates are from individual plants. For metabolite analysis leaf material was harvested 5 days after agroinfiltration by flash freezing in tubes containing metal beads.

### Split-Luciferase assays in *N. benthamiana*

Leaf disks (*ca*. 1 cm) were punched from four biological replicates three days post-infiltration and placed into a custom-made high-density polyethylene (HDPE) multi-well plate. The abaxial side of the leaf was facing up. 200 µl of 0.5 mM d-luciferin (Promega) was added to the leaf disks in each well. The plate was imaged with a NightSHADE LB 985 (Berthold Technologies) with luminescence emission at 0.1 s (wavelength filter 650 nm, 10% intensity) and 8 x 8 pixel binning. Images were taken after 15 and 20 min of incubation with luciferin in the dark with 10s and 2s exposure times, respectively. The pictures were analyzed with indiGOTM 1.4 software (Berthold Technologies) and a scale from 5000-65,000 cps was applied.

### Virus induced gene silencing (VIGS) in *C. roseus*

VIGS was performed as previously described (Li *et al*., 2023b). Briefly, 300 bp target regions of the coding regions of *C. roseus ICYC* and *ISY* genes (transcripts CRO_01G006740.1 and CRO_07G007680.1 in https://datadryad.org/dataset/doi:10.5061/dryad.d2547d851, therein https://datadryad.org/downloads/file_stream/2121644) were selected using the SGN VIGS tool to avoid off-target gene silencing (https://vigs.solgenomics.net/) (Fernandez-Pozo *et al*, 2015). Genomic DNA was extracted from *C. roseus* leaves with the DNeasy Plant Mini Kit (Qiagen). Target region fragments were PCR-amplified from gDNA with in-fusion cloning overhangs using Phusion High-Fidelity DNA Polymerase (ThermoFisher) and specific primers (Supplementary Table 7). The obtained PCR fragments were cloned in the BamHI and XhoI (NEB) digested VIGS vector pTRV2-MgChl (Liscombe & O’Connor, 2011) using the In-Fusion Snap Assembly Master Mix (TaKaRa Bio).

*A. tumefaciens* GV3101 was transformed by electroporation as described above. For inoculation, *A. tumefaciens* GV3101 with plasmid pTRV1 (Liu *et al*, 2002) and *A. tumefaciens* GV3101 carrying pTRV2-MgChl, pTRV2-CrICYC or pTRV2-CrISY were grown over night in a rotary shaker at 28 °C and 300 rpm in each 10 ml of LB medium supplemented with 50 mg/l kanamycin, 25 mg/l gentamicin and 100 mg/l rifampicin to an OD_600_ of ∼2. Cultures were centrifuged for 10 min at 3000 rpm and pellets resuspended in infiltration buffer (100 µM acetosyringone, 10 mM NaCl and 1.75 mM CaCl_2_) to an OD_600_ of 2. After 2 h of incubation on a rotary shaker at room temperature and 60 rpm, 450 µl of each bacterial strain containing a pTRV2 plasmid were mixed with the same volume of the bacterial strain with pTRV1. VIGS inoculation was performed by pipetting 10 µl of the mixed bacterial suspension between plant stem and petiole of the first true leaf of a 30-day-old *C. roseus* cultivar ‘Atlantis Burgundy Halo’ plant (grown in a growth chamber at 16/8 hours light/dark 23°C/21°C, 50% humidity). The stem was pierced with a ø 0.40 x 25 mm Sterican® needle (www.bbraun.com) twice through the bacterial suspension drop. Six plants per pTRV2 construct were inoculated. Plants inoculated with the pTRV2-MgChl strain served as controls. After VIGS inoculation, plants were grown under the same conditions as described above. Yellow tissues (due to co-silencing of *MAGNESIUM CHELATASE SUBUNIT H* gene) were harvested three weeks after inoculation.

### qPCR

Total RNA of *C. roseus* silenced tissues was extracted using the RNeasy Plant Mini Kit (Qiagen) according to the manufacturer’s instructions, including on column DNAse digest. RNA concentrations were measured with a Nanophotometer N60 (Implen). mRNA was reverse-transcribed with 0.5 µg total RNA as input using the iscript cDNA synthesis kit (Biorad) according to the manufacturer’s instructions. cDNA was then diluted 1:8 and 2 µl of the dilution were used in each qPCR reaction of 10 µl total volume in a 96-well plate with 333 nM of each primer (Supplementary Table 7) and 5 µl of PowerUp SYBR Green Master mix (Applied Biosystems) according to manufacturer’s instructions. Amplification was done on a QuantStudio 1 (Applied Biosystems) qPCR machine with the following cycling conditions: 50°C for 2 min, 95°C for 2 min, followed by 40 cycles of 95°C for 1 s and 60°C for 30 s (ramp rate was 1.6°C/s in all cases). Amplification was followed by a melting curve (0.15°C/s from 60°C to 95°C) to confirm the presence of single amplification products. Data was analyzed using the ΔΔCt method and normalized to the expression of the established reference gene *N2227* (Pollier *et al*, 2014). Graphs and statistics were done in Prism Graphpad 10.4.1.

### Metabolite extraction

Leaf material of agroinfiltrated *N. benthamiana* or silenced *C. roseus* was ground using two 4-mm metal beads and a TissueLyser (Qiagen) with pre-cooled adapters, and extracted with 30 µl per mg of fresh material of 70 % MeOH containing 0.1 % formic acid and 1 µM harpagoside as internal standard. Samples were sonicated for 10 min, incubated on a rotator for 15 min and centrifuged at 18000 *g* for 15 min. The supernatants were filtered through a 0.45 µm low binding hydrophilic PTFE filter plate (MultiScreen Solvinert 96, Merck-Millipore) into a 96-well Microtiter Plate (SureSTART WebSeal, Thermo Scientific) according to manufacturer’s instructions. Plates were sealed with Rapid Slit Seal (BioChromato) and immediately analyzed with UPLC-MS/MS.

### Detection of iridoid glucosides by UPLC-MS/MS

The system consisted of an UltiMate 3000 Ultra-High Performance Liquid Chromatography (UHPLC) system (Thermo Fisher Scientific) coupled to an Impact II high resolution Quadrupole Time-Of-Flight (Q-TOF) mass spectrometer (Bruker Daltonics). A Kinetex XB C18 (2.1 x 100 mm, 2.6 µm; 100 Å) column (Phenomenex) was set at 40°C and 0.6 ml/min flow rate and 2 µl of samples were injected. The mobile phase was A:B where A was water with 0.1% formic acid and B was acetonitrile. The gradient was as follows: 5% B at 0.5 min to 30% B at 6 min. Then the column was flushed at 100% B until 7.6 min and re-equilibrated to 5% B until 10 min. Ionization was performed in negative electrospray ionization mode (ESI–) with 3500 V capillary voltage and 500 V end plate offset; a nebulizer pressure of 2.5 bar, with nitrogen at 250°C and a flow of 11 l/min as the drying gas. Acquisition was done at 12 Hz following a mass range from 100 – 1000 *m/z* with data dependent MS/MS and an active exclusion window of 0.2 min, a reconsideration threshold of 1.8-fold change. Fragmentation was triggered on an absolute threshold of 400 and limited to a total cycle time range of 0.5 s. For collision energy, the stepping option model (from 20 to 50 eV) was used. Recalibration of the *m/z* values took place at the start of each run using the expected cluster ion *m/z* values of a direct source infusion of sodium formate-isopropanol solution. At the first minute of each run the LC input was redirected to waste. During this time, the m/z values of the instrument were calibrated using the cluster ion m/z values of a sodium formate-isopropanol solution injected by direct source infusion with a 5 ml syringe connected to an external pump at a flow rate of 0.18 ml/h.

### UPLC-MS/MS data analysis

Raw data was converted to mzml format using MSConvert 3.0.25036-69e37b6 and imported to MZmine 4.5.37 (Chambers *et al*, 2012; Schmid *et al*, 2023). Extracted ion chromatogram (EIC) traces and MS^2^ data of compounds of interest were exported from MZmine. Peak areas were calculated using the MZmine Processing Wizard and exported. Peak areas of compounds of interest were normalized to the internal standard harpagoside and converted to intensity per second. Further data analysis and construction of graphs was done in GraphPad Prism 10.4.1 for Mac OS X.

### Commercially available chemicals and standards

Secologanin (50741), loganic acid (PHL80492), 4-nitrophenyl acetate (N8130) and D-Camphor (50843) were obtained from Sigma. NADPH tetrasodium salt (10621692001) was obtained from Roche. 8-oxogeranial (D476180) and 7*S*-*cis-trans* nepetalactol (N390065) were purchased from Toronto Research Chemicals (TRC). Harpagoside (7471.2) was purchased from Roth.

### Synthesized standards and substrates

Secologanic acid standard was produced as previously described through alkaline hydrolysis of secologanin (Chapelle, 1976; Colinas *et al*., 2025). Secologanin was incubated with 0.1 M NaOH (40 µl per 1 mg secologanin) for five hours and then neutralized with HCl. The completeness of the reaction was confirmed through LC-MS analysis and the solution stored at –25°C. *S*-8-oxocitronellal, *R*-8-oxocitronellal and 7*R*-*cis-cis* nepetalactone were obtained as previously described (Geu-Flores *et al*., 2012; Hernandez Lozada *et al*., 2022; Lichman *et al*., 2019a).

### Recombinant protein production and purification

Expression and purification were performed as previously described with modifications (Stavrinides *et al*, 2016). Briefly, *E. coli* SoluBL21 (DE3) were transformed by heat shock with pOPINF constructs. Pre-cultures were inoculated from single colonies, grown over night at 37°C and used to inoculate 100 ml 2 x YT medium (500 ml in the case of CiICYC_D210A mutant). Cultures were grown at 37°C until OD_600_ 0.5-0.6 was reached, cooled to room temperature and induced with 0.2 mM Isopropyl β-D-1-thiogalactopyranoside (IPTG). After induction the cultures were shifted to 18°C over-night and harvested the next day by centrifugation. Cell pellets were lysed on ice for 30 min with approximately 6 ml lysis buffer per 1 g of cell pellet (50 mM Tris-HCl pH8, 50 mM glycine, 5% glycerol, 500 mM NaCl, 20 mM imidazole, 0.2 mg/ml lysozyme, 1 tablet / 50 ml of complete EDTA free protease inhibitor (Roche)) and sonicated for 2.5 min (2 s on, 3 s off) on ice (Bandelin UW 2070). Supernatants were incubated with gentle shaking in falcon tubes with 250 µl Ni-NTA Agarose (Qiagen) for one hour at 4°C to allow binding of His-tagged proteins. Slurry was pelleted gently by centrifugation at 1000 g for 30 s. The supernatant was removed and the slurry was washed three times with ice cold wash buffer (50 mM Tris-HCL pH8, 50 mM glycine, 5% glycerol, 500 mM NaCl, 20 mM imidazole) by inversion, centrifugation and removal of supernatant. Proteins were eluted using elution buffer (as wash buffer but containing 500 mM imidazole). Elution fractions were concentrated and buffer was exchanged to storage buffer (20 mM HEPES, 150 mM NaCl, pH 7.5) using Amicon Ultra Centrifugal Filters (Millipore) with 10 kDa molecular weight cutoff according to manufacturer’s instructions. Purity was assessed through SDS-PAGE and concentration was determined using the extinction coefficient and measuring the absorbance at 280 nm. Proteins were flash frozen in small aliquots in liquid nitrogen and stored at –70°C.

### In vitro enzyme assays

With 8-oxogeranial as a substrate, enzymes were assayed under the following conditions: 100 mM MOPS pH 7.5, 0.5 mM 8-oxogeranial, 1 mM NADPH, 0.25 µM CrISY or 0.5 µM AmISY, and 2 µM ICYC, NmMLPL or bovine serum album (BSA) as control as indicated. The reactions were set up in a 100 µl total volume and started by the addition of NADPH. Reactions were incubated for 3h at 30°C and 400 rpm. For sequential reactions (Supplementary Fig. 15) ICYC, NmMLPL or BSA were added after 1.5 hours of incubation with CrISY and incubation was continued for additional 1.5 hours. For assays with *S*-, or *R*-8-oxocitronellal, conditions were as follows: 500 mM MOPS pH7.5, ca. 0.5 mM *S*-8-oxocitronellal or ca. 0.1 mM *R*-8-oxocitronellal, respectively, 5 µM of ICYC, NmMLPL or BSA. Reactions were incubated for 16 hours at 30°C and 400 rpm. After incubation camphor (5 or 10 µl of a 1 mM solution in acetonitrile) was added as internal standard to each reaction and briefly vortexed. Reactions were then extracted with 200 µl ethyl acetate and vortexed thoroughly for 2 min. Layers were separated by centrifuging at 18000g for 5 min and 100 µl of the upper ethyl acetate layer was transferred to a glass vial. Samples were immediately analyzed by GC-MS.

Esterase activities were measured in a spectrophotometric assay with 4-nitrophenyl acetate (Hicks *et al*, 2011). Reactions were performed in 96-well plates (CytoOne) in 250 µl total volume. Reactions contained 100 mM HEPES pH 7, 1 mM CaCl_2_, 2.6 mM NaCl, 2 mM 4-nitrophenyl acetate and 0.5 µM of enzyme or BSA as control. The reactions were started by the addition of 4-nitrophenyl acetate through a multi-channel pipette and the plate was immediately placed in a CLARIOstar Plus microplate reader (BMG Labtech). Absorbance at 405 nm was recorded every minute for a total duration of 60 min at 25°C. Data was exported from CLARIOstar Data analysis software (MARS 3.4.1) and further analyzed in Prism GraphPad 10.4.1.

### Detection of iridoids by GC-MS

Samples were analyzed on an achiral column using an Agilent 8890GC system, an Agilent 5977B GC/MSD detector, and a CTC Analytics PAL RSI 120 autosampler system. A Zebron ZB-5^TM^ Plus column (Phenomenex; I.D. = 0.25 mm; L = 30 m; film thickness = 0.25 µm) was used for seperation. 1 µl of samples were injected at 230°C inlet temperature. The carrier gas was helium at 1.1 ml/min constant flow. The temperature gradient was as follows: 2 min at 60°C, ramp up to 220°C at 7 °C/min, ramp up to 300°C at 60°C/min, 2 min at 300°C. The MSD transfer line temperature was 280°C and the MS source temperature 230°C. After a solvent delay of 6 min, a mass range of 35-250 amu was collected at 70 eV fragmentation energy.

For confirmation of 7*R*-*cis-cis* nepetalactone identity, samples were analyzed on a chiral column using previously established conditions (Hernandez Lozada *et al*., 2022). Briefly, samples were analyzed on an Agilent system consisting of an 8890GC system, a 5977B GC/MSD detector, and a 7693 A autosampler. A Supelco β-DEX^TM^ 225 column (I.D. = 0.25 mm; L = 30 m; film thickness = 0.25 µm) was used for separation. The samples (1 µl) were injected at 220 °C inlet temperature using a split ratio of 1:10. Helium was used as carrier gas at a flow rate of 1.1 ml/min. The oven temperature ramp was as follows: 3 min at 80 °C, ramp up to 120°C at 10 °C/min, 45 min at 120°C, ramp up to 200°C at 10 °C/min, 2 min at 200°C. The MSD transfer line temperature was 220°C and the MS source at 230°C. After 10 min of solvent delay, a mass range of 50-350 amu was collected at 70eV fragmentation energy.

### Synthesis of 7*R*-*cis-cis* nepetalactone from enzymatically produced 7*R*-*cis-cis* nepetalactol

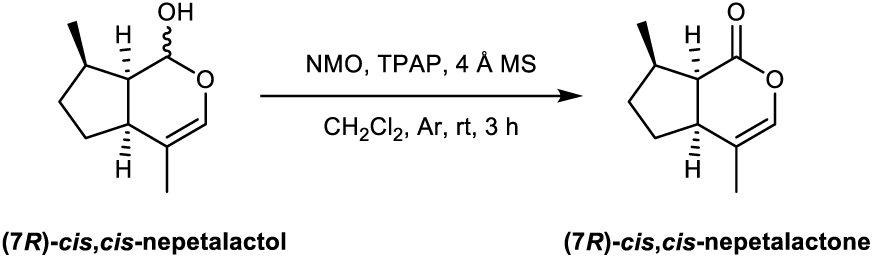

The combined assays (5x 100 µl reactions with AmISY and AmICYC performed as described above) were extracted with CH_2_Cl_2_ (100 µl x 3) and the resulting organic phase was then dried over anhydrous Na_2_SO_4_ and partially concentrated under a gentle stream of Ar. To the concentrated organic extract was then sequentially added 4 Å MS (*ca*. 5 mg), *N*-methylmorpholine *N*-oxide (1 mg, 8.54 µmol), and tetra-*n*-propylammonium perruthenate (1 mg, 2.85 µmol). After stirring under an Ar atmosphere at room temperature for 3 h, the reaction mass was filtered through a short pad of Celite^®^ and then directly passed through a short silica column, eluting with CH_2_Cl_2_ (2 ml) and Et_2_O (2 ml), to give a colorless eluate that was then partially concentrated under a gentle stream of Ar and directly submitted for chiral GC-MS analysis.

### GC-MS data analysis

Data was analyzed in Agilent MassHunter Qualitative Analysis 10.0. Total ion chromatograms (TICs) and were exported as CSV files and imported to Prism Graphpad version 10.4.1 for visualization. TICs shown in the same graph are with the same scaled Y-axis unless otherwise indicated. Peak areas were calculated in Agilent MassHunter Qualitative Analysis 10.0, exported and normalized to internal standard camphor. Bar graphs were constructed in Prism Graphpad version 10.4.1.

### Phylogenetic analyses

Sequences of ICYC orthologs and other methylesterases were obtained through blast searches against TAIR11, NCBI and 1KP (One Thousand Plant Transcriptomes, 2019) databases, against transcriptome assemblies from the MintGenomics project (Mint Evolutionary Genomics Consortium. Electronic address & Mint Evolutionary Genomics, 2018), or against genome data obtained during this study. Sequences and accession numbers are provided in Supplementary dataset 1. Full-length amino acid sequences were aligned with webPRANK (https://www.ebi.ac.uk/goldman-srv/webprank/) (Löytynoja & Goldman, 2010). Sequence logos were obtained by importing alignments to Geneious Prime 2025.1.2 (Dotmatics). The IQ-TREE webserver (http://iqtree.cibiv.univie.ac.at/) was used to build Maximum Likelihood phylogenetic trees (automatic substitution model; bootstrap value 1000) (Trifinopoulos *et al*, 2016). Trees were visualized in iTOL and graphically edited using itol (https://itol.embl.de/) (Letunic & Bork, 2024) and Adobe Illustrator 27.8.

### Protein models and Docking

Protein models were predicted by AlphaFold3 through the AlphaFold Server (https://alphafoldserver.com/) (Abramson *et al*, 2024). Docking was done using Autodock Vina on the SwissDock webserver (https://www.swissdock.ch/) (Bugnon *et al*, 2024; Eberhardt *et al*, 2021). Models were visualized using ChimeraX version 1.8 for Mac (Meng *et al*, 2023).

## References

Abramson J, Adler J, Dunger J, Evans R, Green T, Pritzel A, Ronneberger O, Willmore L, Ballard AJ, Bambrick J et al (2024) Accurate structure prediction of biomolecular interactions with AlphaFold 3. Nature 630: 493–500

Afgan E, Baker D, Batut B, van den Beek M, Bouvier D, Cech M, Chilton J, Clements D, Coraor N, Gruning BA et al (2018) The Galaxy platform for accessible, reproducible and collaborative biomedical analyses: 2018 update. Nucleic Acids Res 46: W537–W544

Albach DC, Soltis PS, Soltis DE (2001) Patterns of Embryological and Biochemical Evolution in the Asterids. Systematic Botany 26: 242–262

Altschul SF, Gish W, Miller W, Meyers EW, Lipman DJ (1990) Basic Local Alignment Search Tool. J Mol Biol 215: 403–410

Awadasseid A, Li W, Liu Z, Qiao C, Pang J, Zhang G, Luo Y (2020) Characterization of Camptotheca acuminata 10-hydroxygeraniol oxidoreductase and iridoid synthase and their application in biological preparation of nepetalactol in Escherichia coli featuring NADP(+) - NADPH cofactors recycling. Int J Biol Macromol 162: 1076–1085

Berrow NS, Alderton D, Sainsbury S, Nettleship J, Assenberg R, Rahman N, Stuart DI, Owens RJ (2007) A versatile ligation-independent cloning method suitable for high-throughput expression screening applications. Nucleic Acids Res 35: e45

Birkett MA, Hassanali A, Hoglund S, Pettersson J, Pickett JA (2011) Repellent activity of catmint, Nepeta cataria, and iridoid nepetalactone isomers against Afro-tropical mosquitoes, ixodid ticks and red poultry mites. Phytochemistry 72: 109–114

Bolger AM, Lohse M, Usadel B (2014) Trimmomatic: a flexible trimmer for Illumina sequence data. Bioinformatics 30: 2114–2120

Brůna T, Hoff KJ, Lomsadze A, Stanke M, Borodovsky M (2021) BRAKER2: automatic eukaryotic genome annotation with GeneMark-EP+ and AUGUSTUS supported by a protein database. NAR Genom Bioinform 3: lqaa108

Bugnon M, Rohrig UF, Goullieux M, Perez MAS, Daina A, Michielin O, Zoete V (2024) SwissDock 2024: major enhancements for small-molecule docking with Attracting Cavities and AutoDock Vina. Nucleic Acids Res 52: W324–W332

Burlat V, Oudin A, Courtois M, Rideau M, St-Pierre B (2004) Co-expression of three MEP pathway genes and geraniol 10-hydroxylase in internal phloem parenchyma of Catharanthus roseus implicates multicellular translocation of intermediates during the biosynthesis of monoterpene indole alkaloids and isoprenoid-derived primary metabolites. Plant J 38: 131–141

Burse A, Boland W (2017) Deciphering the route to cyclic monoterpenes in Chrysomelina leaf beetles: source of new biocatalysts for industrial application? Z Naturforsch C J Biosci 72: 417–427

Campbell MS, Law M, Holt C, Stein JC, Moghe GD, Hufnagel DE, Lei J, Achawanantakun R, Jiao D, Lawrence CJ et al (2014) MAKER-P: a tool kit for the rapid creation, management, and quality control of plant genome annotations. Plant Physiol 164: 513–524

Cardenas PD, Sonawane PD, Heinig U, Jozwiak A, Panda S, Abebie B, Kazachkova Y, Pliner M, Unger T, Wolf D et al (2019) Pathways to defense metabolites and evading fruit bitterness in genus Solanum evolved through 2-oxoglutarate-dependent dioxygenases. Nat Commun 10: 5169

Carrier G, Santoni S, Rodier-Goud M, Canaguier A, Kochko A, Dubreuil-Tranchant C, This P, Boursiquot JM, Le Cunff L (2011) An efficient and rapid protocol for plant nuclear DNA preparation suitable for next generation sequencing methods. Am J Bot 98: e13–15

Chambers MC, Maclean B, Burke R, Amodei D, Ruderman DL, Neumann S, Gatto L, Fischer B, Pratt B, Egertson J et al (2012) A cross-platform toolkit for mass spectrometry and proteomics. Nat Biotechnol 30: 918–920

Chanderbali AS, Dervinis C, Anghel IG, Soltis DE, Soltis PS, Zapata F (2024) Draft genome assemblies for two species of Escallonia (Escalloniales). BMC Genomic Data 25: 1

Chapelle JP (1976) VOGELOSIDE ET ACIDE SECOLOGANIQUE, GLUCOSIDES SECOIRIDOIDES D’ANTHOCLEISTA VOGELU. Planta Med 29: 268–274

Chen H, Zou Y, Shang Y, Lin H, Wang Y, Cai R, Tang X, Zhou JM (2008) Firefly luciferase complementation imaging assay for protein-protein interactions in plants. Plant Physiol 146: 368–376

Chen N (2004) Using RepeatMasker to identify repetitive elements in genomic sequences. Curr Protoc Bioinformatics Chapter 4: Unit 4 10

Colinas M, Morweiser C, Dittberner O, Chioca B, Alam R, Leucke H, Nakamura Y, Guerrero DAS, Heinicke S, Kunert M et al (2024) Independent evolution of ipecac alkaloid biosynthesis. bioRxiv: 2024.2009.2023.614470

Colinas M, Morweiser C, Dittberner O, Chioca B, Alam R, Leucke H, Nakamura Y, Serna Guerrero DA, Heinicke S, Kunert M et al (2025) Ipecac alkaloid biosynthesis in two evolutionarily distant plants. Nat Chem Biol

Colinas M, Pollier J, Vaneechoutte D, Malat DG, Schweizer F, De Milde L, De Clercq R, Guedes JG, Martinez-Cortes T, Molina-Hidalgo FJ et al (2021) Subfunctionalization of Paralog Transcription Factors Contributes to Regulation of Alkaloid Pathway Branch Choice in Catharanthus roseus. Front Plant Sci 12: 687406

Collu G, Unver N, Peltenburg-Looman AM, van der Heijden R, Verpoorte R, Memelink J (2001) Geraniol 10-hydroxylase, a cytochrome P450 enzyme involved in terpenoid indole alkaloid biosynthesis. FEBS Lett 508: 215–220

Dinda B, Debnath S, Harigaya Y (2007) Naturally Occurring Secoiridoids and Bioactivity of Naturally Occurring Iridoids and Secoiridoids. A Review, Part 2. Chemical and Pharmaceutical Bulletin 55: 689–728

Dobler S, Petschenka G, Pankoke H (2011) Coping with toxic plant compounds--the insect’s perspective on iridoid glycosides and cardenolides. Phytochemistry 72: 1593–1604

Dogru E, Warzecha H, Seibel F, Haebel S, Lottspeich F, Stockigt J (2000) The gene encoding polyneuridine aldehyde esterase of monoterpenoid indole alkaloid biosynthesis in plants is an ortholog of the alpha/betahydrolase super family. Eur J Biochem 267: 1397–1406

Dudley QM, Jo S, Guerrero DAS, Chhetry M, Smedley MA, Harwood WA, Sherden NH, O’Connor SE, Caputi L, Patron NJ (2022) Reconstitution of monoterpene indole alkaloid biosynthesis in genome engineered Nicotiana benthamiana. Commun Biol 5: 949

Eberhardt J, Santos-Martins D, Tillack AF, Forli S (2021) AutoDock Vina 1.2.0: New Docking Methods, Expanded Force Field, and Python Bindings. J Chem Inf Model 61: 3891–3898

Eisner T (1964) Catnip: Its *Raison d’Etre*. Science 146: 1318–1320

Farrow SC, Kamileen MO, Caputi L, Bussey K, Mundy JEA, McAtee RC, Stephenson CRJ, O’Connor SE (2019) Biosynthesis of an Anti-Addiction Agent from the Iboga Plant. J Am Chem Soc 141: 12979–12983

Fernandez-Pozo N, Rosli HG, Martin GB, Mueller LA (2015) The SGN VIGS tool: user-friendly software to design virus-induced gene silencing (VIGS) constructs for functional genomics. Mol Plant 8: 486–488

Flynn JM, Hubley R, Goubert C, Rosen J, Clark AG, Feschotte C, Smit AF (2020) RepeatModeler2 for automated genomic discovery of transposable element families. Proc Natl Acad Sci U S A 117: 9451–9457

Forouhar F, Yang Y, Kumar D, Chen Y, Fridman E, Park SW, Chiang Y, Acton TB, Montelione GT, Pichersky E et al (2005) Structural and biochemical studies identify tobacco SABP2 as a methyl salicylate esterase and implicate it in plant innate immunity. Proceedings of the National Academy of Sciences 102: 1773–1778

Fuji Y, Uchida A, Fukahori K, Chino M, Ohtsuki T, Matsufuji H (2018) Chemical characterization and biological activity in young sesame leaves (Sesamum indicum L.) and changes in iridoid and polyphenol content at different growth stages. PLoS One 13: e0194449

Geu-Flores F, Sherden NH, Courdavault V, Burlat V, Glenn WS, Wu C, Nims E, Cui Y, O’Connor SE (2012) An alternative route to cyclic terpenes by reductive cyclization in iridoid biosynthesis. Nature 492: 138–142

Guan D, McCarthy SA, Wood J, Howe K, Wang Y, Durbin R (2020) Identifying and removing haplotypic duplication in primary genome assemblies. Bioinformatics 36: 2896–2898

Haas BJ, Delcher AL, Mount SM, Wortman JR, Smith RK, Jr., Hannick LI, Maiti R, Ronning CM, Rusch DB, Town CD et al (2003) Improving the Arabidopsis genome annotation using maximal transcript alignment assemblies. Nucleic Acids Res 31: 5654–5666

Hernandez Lozada NJ, Hong B, Wood JC, Caputi L, Basquin J, Chuang L, Kunert M, Rodriguez Lopez CE, Langley C, Zhao D et al (2022) Biocatalytic routes to stereo-divergent iridoids. Nat Commun 13: 4718

Hicks MA, Barber AE, 2nd, Giddings LA, Caldwell J, O’Connor SE, Babbitt PC (2011) The evolution of function in strictosidine synthase-like proteins. Proteins 79: 3082–3098

Hong B, Grzech D, Caputi L, Sonawane P, Lopez CER, Kamileen MO, Hernandez Lozada NJ, Grabe V, O’Connor SE (2022) Biosynthesis of strychnine. Nature 607: 617–622

Irmler S, Schröder G, St-Pierre B, Crouch NP, Hotze M, Schmidt J, Strack D, Matern U, Schröder J (2008) Indole alkaloid biosynthesis in Catharanthus roseus: new enzyme activities and identification of cytochrome P450 CYP72A1 as secologanin synthase. The Plant Journal 24: 797–804

Jensen S (1991) Plant iridoids, their Biosynthesis and Distribution in Angiosperms. In: pp. 133–158.

Jensen SR (1992) Systematic Implications of the Distribution of Iridoids and Other Chemical Compounds in the Loganiaceae and Other Families of the Asteridae. Annals of the Missouri Botanical Garden 79: 284–302

Jensen SR, Nielsen BJ (1980) Iridoid glucosides in Griselinia, Aralidium and Toricellia. Phytochemistry 19: 2685–2688

Kakuda R, Imai M, Yaoita Y, Machida K, Kikuchi M (2000) Secoiridoid glycosides from the flower buds of Lonicera japonica. Phytochemistry 55: 879–881

Kaminow B, Yunusov D, Dobin A (2021) STARsolo: accurate, fast and versatile mapping/quantification of single-cell and single-nucleus RNA-seq data. bioRxiv

Kim D, Paggi JM, Park C, Bennett C, Salzberg SL (2019) Graph-based genome alignment and genotyping with HISAT2 and HISAT-genotype. Nat Biotechnol 37: 907–915

Kollner TG, David A, Luck K, Beran F, Kunert G, Zhou JJ, Caputi L, O’Connor SE (2022) Biosynthesis of iridoid sex pheromones in aphids. Proc Natl Acad Sci U S A 119: e2211254119

Kolmogorov M, Yuan J, Lin Y, Pevzner PA (2019) Assembly of long, error-prone reads using repeat graphs. Nat Biotechnol 37: 540–546

Koudounas K, Banilas G, Michaelidis C, Demoliou C, Rigas S, Hatzopoulos P (2015) A defence-related Olea europaea beta-glucosidase hydrolyses and activates oleuropein into a potent protein cross-linking agent. J Exp Bot 66: 2093–2106

Kovaka S, Zimin AV, Pertea GM, Razaghi R, Salzberg SL, Pertea M (2019) Transcriptome assembly from long-read RNA-seq alignments with StringTie2. Genome Biol 20: 278

Kries H, Kellner F, Kamileen MO, O’Connor SE (2017) Inverted stereocontrol of iridoid synthase in snapdragon. J Biol Chem 292: 14659–14667

Lamesch P, Berardini TZ, Li D, Swarbreck D, Wilks C, Sasidharan R, Muller R, Dreher K, Alexander DL, Garcia-Hernandez M et al (2012) The Arabidopsis Information Resource (TAIR): improved gene annotation and new tools. Nucleic Acids Res 40: D1202–1210

Lawas LMF, Kamileen MO, Buell CR, O’Connor SE, Leisner CP (2023) Transcriptome-based identification and functional characterization of iridoid synthase involved in monotropein biosynthesis in blueberry. Plant Direct 7: e512

Lemenager D, Ouelhazi L, Mahroug S, Veau B, St-Pierre B, Rideau M, Aguirreolea J, Burlat V, Clastre M (2005) Purification, molecular cloning, and cell-specific gene expression of the alkaloid-accumulation associated protein CrPS in Catharanthus roseus. J Exp Bot 56: 1221–1228

Letunic I, Bork P (2024) Interactive Tree of Life (iTOL) v6: recent updates to the phylogenetic tree display and annotation tool. Nucleic Acids Res 52: W78–W82

Li C, Colinas M, Wood JC, Vaillancourt B, Hamilton JP, Jones SL, Caputi L, O’Connor SE, Buell CR (2025) Cell-type-aware regulatory landscapes governing monoterpene indole alkaloid biosynthesis in the medicinal plant Catharanthus roseus. New Phytol 245: 347–362

Li C, Deans NC, Buell CR (2023a) “Simple Tidy GeneCoEx”: A gene co-expression analysis workflow powered by tidyverse and graph-based clustering in R. Plant Genome 16: e20323

Li C, Wood JC, Deans NC, Jarrell AF, Martin D, Mailloux K, Wang Y-W, Robin Buell C (2022) Nuclei isolation protocol from diverse angiosperm species. bioRxiv

Li C, Wood JC, Vu AH, Hamilton JP, Rodriguez Lopez CE, Payne RME, Serna Guerrero DA, Gase K, Yamamoto K, Vaillancourt B et al (2023b) Single-cell multi-omics in the medicinal plant Catharanthus roseus. Nat Chem Biol 19: 1031–1041

Li H (2021) New strategies to improve minimap2 alignment accuracy. Bioinformatics 37: 4572–4574

Li M, Zhang D, Gao Q, Luo Y, Zhang H, Ma B, Chen C, Whibley A, Zhang Y, Cao Y et al (2019) Genome structure and evolution of Antirrhinum majus L. Nat Plants 5: 174–183

Li W, Cowley A, Uludag M, Gur T, McWilliam H, Squizzato S, Park YM, Buso N, Lopez R (2015) The EMBL-EBI bioinformatics web and programmatic tools framework. Nucleic Acids Res 43: W580–584

Li Y, Wei H, Yang J, Du K, Li J, Zhang Y, Qiu T, Liu Z, Ren Y, Song L et al (2020) High-quality de novo assembly of the Eucommia ulmoides haploid genome provides new insights into evolution and rubber biosynthesis. Hortic Res 7: 183

Lichman BR (2021) The scaffold-forming steps of plant alkaloid biosynthesis. Nat Prod Rep 38: 103–129

Lichman BR, Godden GT, Hamilton JP, Palmer L, Kamileen MO, Zhao D, Vaillancourt B, Wood JC, Sun M, Kinser TJ et al (2020) The evolutionary origins of the cat attractant nepetalactone in catnip. Science Advances 6: eaba0721

Lichman BR, Kamileen MO, Titchiner GR, Saalbach G, Stevenson CEM, Lawson DM, O’Connor SE (2019a) Uncoupled activation and cyclization in catmint reductive terpenoid biosynthesis. Nat Chem Biol 15: 71–79

Lichman BR, O’Connor SE, Kries H (2019b) Biocatalytic Strategies towards [4+2] Cycloadditions. Chemistry 25: 6864–6877

Lindner S, Geu-Flores F, Brase S, Sherden NH, O’Connor SE (2014) Conversion of substrate analogs suggests a Michael cyclization in iridoid biosynthesis. Chem Biol 21: 1452–1456

Liscombe DK, O’Connor SE (2011) A virus-induced gene silencing approach to understanding alkaloid metabolism in Catharanthus roseus. Phytochemistry 72: 1969–1977

Liu Y, Schiff M, Marathe R, Dinesh-Kumar SP (2002) Tobacco Rar1, EDS1 and NPR1/NIM1 like genes are required for N-mediated resistance to tobacco mosaic virus. Plant J 30: 415–429

Löytynoja A, Goldman N (2010) webPRANK: a phylogeny-aware multiple sequence aligner with interactive alignment browser. BMC Bioinformatics 11: 579

Martin M (2011) Cutadapt removes adapter sequences from high-throughput sequencing reads. EMBnetjournal 17: 10–12

Mattern-Dogru E, Ma X, Hartmann J, Decker H, Stockigt J (2002) Potential active-site residues in polyneuridine aldehyde esterase, a central enzyme of indole alkaloid biosynthesis, by modelling and site-directed mutagenesis. Eur J Biochem 269: 2889–2896

McGinnis CS, Murrow LM, Gartner ZJ (2019) DoubletFinder: Doublet detection in single-cell RNA sequencing data using artificial nearest neighbors. Cell Syst 8: 329–337.e324

Meng EC, Goddard TD, Pettersen EF, Couch GS, Pearson ZJ, Morris JH, Ferrin TE (2023) UCSF ChimeraX: Tools for structure building and analysis. Protein Sci 32: e4792

Miettinen K, Dong L, Navrot N, Schneider T, Burlat V, Pollier J, Woittiez L, van der Krol S, Lugan R, Ilc T et al (2014) The seco-iridoid pathway from Catharanthus roseus. Nat Commun 5: 3606

Miller JC, Schuler MA (2022) Single mutations toggle the substrate selectivity of multifunctional Camptotheca secologanic acid synthases. J Biol Chem 298: 102237

Mint Evolutionary Genomics Consortium. Electronic address bme, Mint Evolutionary Genomics C (2018) Phylogenomic Mining of the Mints Reveals Multiple Mechanisms Contributing to the Evolution of Chemical Diversity in Lamiaceae. Mol Plant 11: 1084–1096

Mistry J, Chuguransky S, Williams L, Qureshi M, Salazar GA, Sonnhammer ELL, Tosatto SCE, Paladin L, Raj S, Richardson LJ et al (2021) Pfam: The protein families database in 2021. Nucleic Acids Res 49: D412–D419

Morita H, Takatsu H, Kobayashi Ji (2003) Daphnezomines P, Q, R and S, new alkaloids from Daphniphyllum humile. Tetrahedron 59: 3575–3579

Murata J, Roepke J, Gordon H, De Luca V (2008) The leaf epidermome of Catharanthus roseus reveals its biochemical specialization. Plant Cell 20: 524–542

Murayama T, Hatakeyama K, Shiraiwa R, Shiono Y, Takahashi K, Ikeda M (2004) A Secoiridoid, 10-Cinnamoyloxyoleoside, Isolated from the Leaves of Helwingia japonica. Natural Medicines 58: 42–45

Ollis DL, Cheah E, Cygler M, Dijkstra B, Frolow F, Franken SM, Harel M, Remington SJ, Silman I, Schrag J et al (1992) The α/β hydrolase fold. Protein Engineering 5: 197–211

One Thousand Plant Transcriptomes I (2019) One thousand plant transcriptomes and the phylogenomics of green plants. Nature 574: 679–685

Pollier J, Vanden Bossche R, Rischer H, Goossens A (2014) Selection and validation of reference genes for transcript normalization in gene expression studies in Catharanthus roseus. Plant Physiol Biochem 83: 20–25

Rauwerdink A, Kazlauskas RJ (2015) How the Same Core Catalytic Machinery Catalyzes 17 Different Reactions: the Serine-Histidine-Aspartate Catalytic Triad of alpha/beta-Hydrolase Fold Enzymes. ACS Catal 5: 6153–6176

Schmid R, Heuckeroth S, Korf A, Smirnov A, Myers O, Dyrlund TS, Bushuiev R, Murray KJ, Hoffmann N, Lu M et al (2023) Integrative analysis of multimodal mass spectrometry data in MZmine 3. Nat Biotechnol 41: 447–449

Simkin AJ, Miettinen K, Claudel P, Burlat V, Guirimand G, Courdavault V, Papon N, Meyer S, Godet S, St-Pierre B et al (2013) Characterization of the plastidial geraniol synthase from Madagascar periwinkle which initiates the monoterpenoid branch of the alkaloid pathway in internal phloem associated parenchyma. Phytochemistry 85: 36–43

Stavrinides A, Tatsis EC, Caputi L, Foureau E, Stevenson CE, Lawson DM, Courdavault V, O’Connor SE (2016) Structural investigation of heteroyohimbine alkaloid synthesis reveals active site elements that control stereoselectivity. Nat Commun 7: 12116

Stull GW, Schori M, Soltis DE, Soltis PS (2018) Character evolution and missing (morphological) data across Asteridae. Am J Bot 105: 470–479

Sun S, Shen X, Li Y, Li Y, Wang S, Li R, Zhang H, Shen G, Guo B, Wei J et al (2023) Single-cell RNA sequencing provides a high-resolution roadmap for understanding the multicellular compartmentation of specialized metabolism. Nat Plants 9: 179–190

Trenti F, Yamamoto K, Hong B, Paetz C, Nakamura Y, O’Connor SE (2021) Early and Late Steps of Quinine Biosynthesis. Org Lett 23: 1793–1797

Trifinopoulos J, Nguyen LT, von Haeseler A, Minh BQ (2016) W-IQ-TREE: a fast online phylogenetic tool for maximum likelihood analysis. Nucleic Acids Res 44: W232–235

Van Moerkercke A, Steensma P, Gariboldi I, Espoz J, Purnama PC, Schweizer F, Miettinen K, Vanden Bossche R, De Clercq R, Memelink J et al (2016) The basic helix-loop-helix transcription factor BIS2 is essential for monoterpenoid indole alkaloid production in the medicinal plant Catharanthus roseus. Plant J 88: 3–12

Van Moerkercke A, Steensma P, Schweizer F, Pollier J, Gariboldi I, Payne R, Vanden Bossche R, Miettinen K, Espoz J, Purnama PC et al (2015) The bHLH transcription factor BIS1 controls the iridoid branch of the monoterpenoid indole alkaloid pathway in *Catharanthus roseus*. Proceedings of the National Academy of Sciences 112: 8130–8135

Viljoen A, Mncwangi N, Vermaak I (2012) Anti-Inflammatory Iridoids of Botanical Origin. Current Medicinal Chemistry 19: 2104–2127

Vlot AC, Liu PP, Cameron RK, Park SW, Yang Y, Kumar D, Zhou F, Padukkavidana T, Gustafsson C, Pichersky E et al (2008) Identification of likely orthologs of tobacco salicylic acid-binding protein 2 and their role in systemic acquired resistance in Arabidopsis thaliana. Plant J 56: 445–456

Volk J, Sarafeddinov A, Unver T, Marx S, Tretzel J, Zotzel J, Warzecha H (2019) Two novel methylesterases from Olea europaea contribute to the catabolism of oleoside-type secoiridoid esters. Planta 250: 2083–2097

Walker BJ, Abeel T, Shea T, Priest M, Abouelliel A, Sakthikumar S, Cuomo CA, Zeng Q, Wortman J, Young SK et al (2014) Pilon: an integrated tool for comprehensive microbial variant detection and genome assembly improvement. PLoS One 9: e112963

Yang Y, Xu R, Ma C-j, Vlot AC, Klessig DF, Pichersky E (2008) Inactive Methyl Indole-3-Acetic Acid Ester Can Be Hydrolyzed and Activated by Several Esterases Belonging to the AtMES Esterase Family of Arabidopsis. Plant Physiology 147: 1034–1045

Yocca AE, Platts A, Alger E, Teresi S, Mengist MF, Benevenuto J, Ferrao LFV, Jacobs M, Babinski M, Magallanes-Lundback M et al (2023) Blueberry and cranberry pangenomes as a resource for future genetic studies and breeding efforts. Hortic Res 10: uhad202

Yu H, Guo K, Lai K, Shah MA, Xu Z, Cui N, Wang H (2022) Chromosome-scale genome assembly of an important medicinal plant honeysuckle. Sci Data 9: 226

Zhang J, Hansen LG, Gudich O, Viehrig K, Lassen LMM, Schrubbers L, Adhikari KB, Rubaszka P, Carrasquer-Alvarez E, Chen L et al (2022) A microbial supply chain for production of the anti-cancer drug vinblastine. Nature 609: 341–347

Zhang Y, Chen M, Siemiatkowska B, Toleco MR, Jing Y, Strotmann V, Zhang J, Stahl Y, Fernie AR (2020) A Highly Efficient Agrobacterium-Mediated Method for Transient Gene Expression and Functional Studies in Multiple Plant Species. Plant Commun 1: 100028

Zuntini AR, Carruthers T, Maurin O, Bailey PC, Leempoel K, Brewer GE, Epitawalage N, Françoso E, Gallego-Paramo B, McGinnie C et al (2024) Phylogenomics and the rise of the angiosperms. Nature

