## Supplementary Figures 1-19 for "Discovery of iridoid cyclase completes the iridoid pathway in asterids"

**Supplementary Information**

### Supplementary Figures

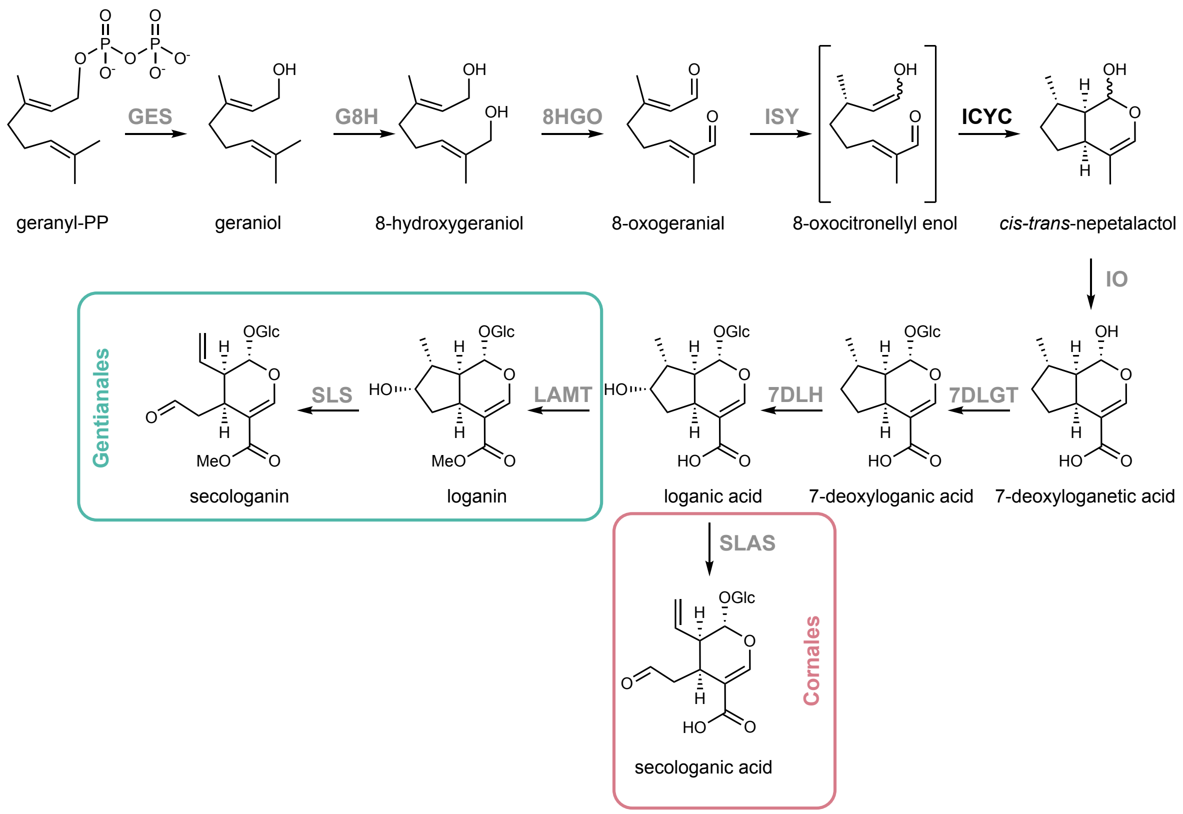

**Supplementary Fig. 1. Secoiridoid pathway in different asterid orders**. Complete secoiridoid pathway. Enzymes labeled in gray have been previously identified in other studies on *Catharanthus roseus* (Gentianales) and/or *Camptotheca acuminata* (Cornales). Generally, in the Gentianales (and other orders) the pathway end product is the methylester secologanin whereas in the Cornales lineage it is secologanic acid due to the absence of LAMT (Kang *et al*, 2021). SLS and SLAS are orthologs and can both accept loganic acid as substrates (Miller *et al*, 2021). GES, geraniol synthase; G8H, geraniol 8-hydroxylase; 8HGO, 8-hydroxygeraniol oxidase; ISY, iridoid synthase; ICYC, iridoid cyclase; IO, iridoid oxidase; 7DLGT, 7-deoxyloganetic acid glucosidase; 7DLH, 7-deoxyloganic acid hydroxylase; SLAS, secologanic acid synthase; LAMT, loganic acid methyltransferase; SLS, secologanin synthase.

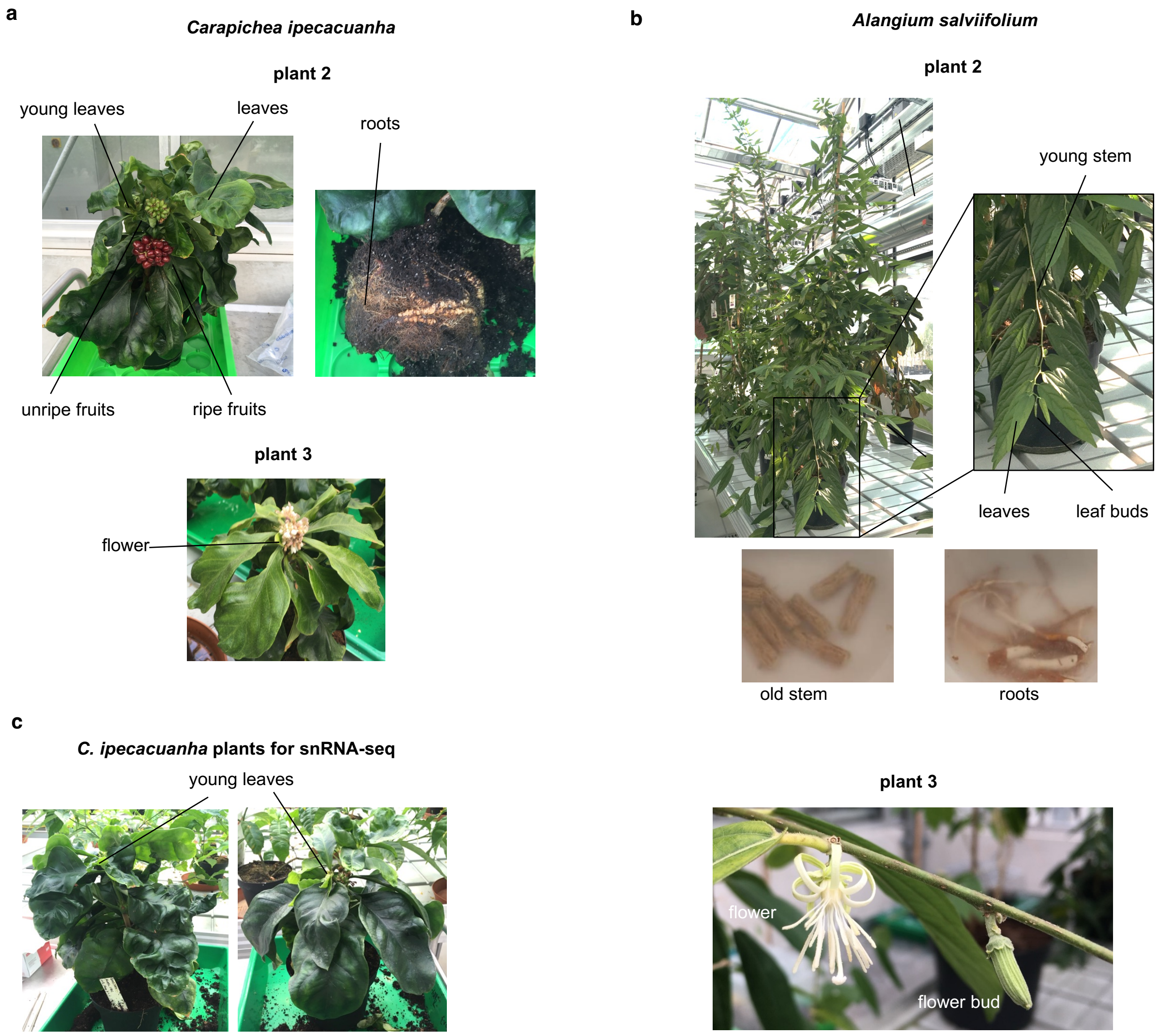

**Supplementary Fig. 2. Photos of plants sampled for genome sequencing, tissue specific RNA-sequencing and single nuclei RNA-sequencing. a**, *Carapichea ipecacuanha* "plant 2" and "plant 3" were 1.5 years old at the time of harvesting. **b**, *Alangium salviifolium* "plant 2" and "plant 3" were 2.5 and 4 years old, respectively at the time of harvesting. **c**, *C. ipecacuanha* young leaves for single nuclei RNA-seq (snRNA-seq) were harvested from 1.5-year-old plants. All plants were grown in a greenhouse with controlled conditions (12/12 hours light/dark 28-30°C/24-26°C, 70-80% humidity). Sequencing data for plants labelled as "plant 1" in this study were published previously (Colinas *et al*, 2025) (https://www.ncbi.nlm.nih.gov/bioproject/PRJNA1169657).

**a**

| enzyme name | short name | Accession *C. roseus* | Best blast hit *C. ipecacuanha* | % identity | Best blast hit *A. salviifolium* | % identity |
|---|---|---|---|---|---|---|
| geraniol synthase | GES | JN882024 | Caipe.S184100 | 79 | AIsal.S224590 | 66.5 |
| geraniol 8-hydroxylase | G8H | KF561461 | Caipe.S125190 | 81 | AIsal.S141850 | 74.2 |
| 8-hydroxygeraniol oxidase | 8HGO | KF302069 | Caipe.S388690 | 87.3 | AIsal.S112090 | 70.1 |
| iridoid synthase | ISY | KJ873886 | Caipe.S317540 | 77.9 | AIsal.S308160 | 61.2 |
| iridoid oxidase | IO | KF591593 | Caipe.S160820 | 89.9 | AIsal.S107160 | 77.8 |
| 7-deoxyloganetic glycosyltransferase | 7DLGT | KF302067 | Caipe.S283170 | 67.1 | AIsal.S289890 | 66 |
| 7-deoxyloganic acid hydroxylase | 7DLH | AB733667 | Caipe.S364920 | 84.6 | AIsal.S217620 | 72.8 |
| loganic acid methyltransferase | LAMT | KF415116 | Caipe.S248290 | 79.6 | AIsal.S158890* | 53 |
| secologanin synthase | SLS | KM524262 | Caipe.S229390 | 76.1 | AIsal.S289930 | 64.3 |

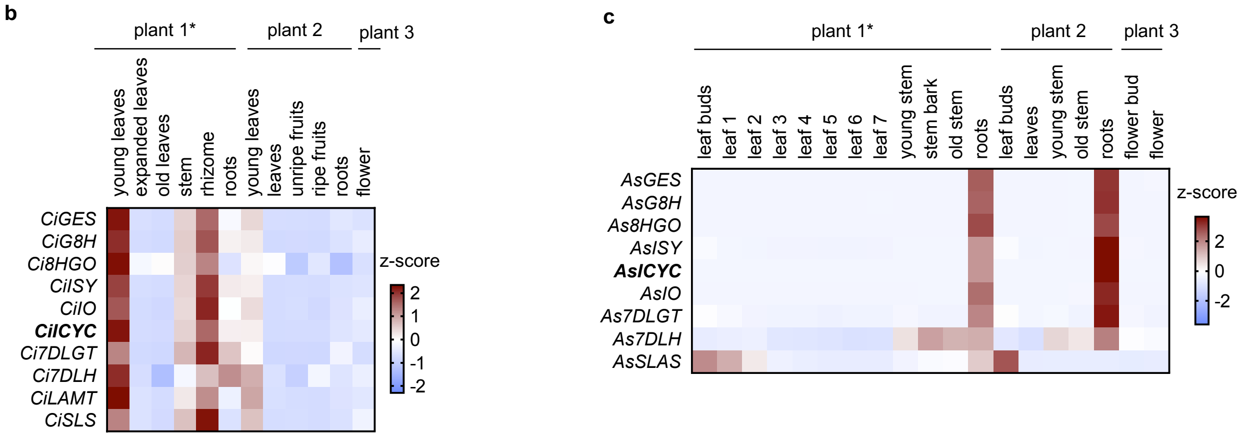

**Supplementary Fig. 3. Identification of orthologs of known *C. roseus* secoiridoid pathway genes in *C. ipecacuanha* and *A. salviifolium* and tissue-specific expression data. a**, Amino acid sequences of the previously published *C. roseus* pathway enzymes were subjected to BLAST (tblastn) against *C. ipecacuanha* and *A. salviifolium* transcripts (working gene models) generated from the annotated genomes. The hits with the highest sequence identities were considered orthologs and the activities of these gene products were confirmed in this study. Note, that for LAMT the hit with the highest sequencing identity in *A. salviifolium* (marked with an asterisk) showed comparably lower sequence identity (Kang *et al.*, 2021). **b,** Tissue-specific expression analysis of identified iridoid pathway gene orthologs and the newly identified *CiICYC* in *C. ipecacuanha* shows high expression of identified iridoid pathway genes ortholog alongside the newly identified *CiICYC* in young leaves and rhizome. **c,** Tissue-specific expression analysis of identified iridoid pathway gene orthologs and the newly identified *AsICYC* in *A. salviifolium* reveals almost exclusive expression in the roots, apart from *AsSLAS*, which is also expressed in leaf buds and young leaves.

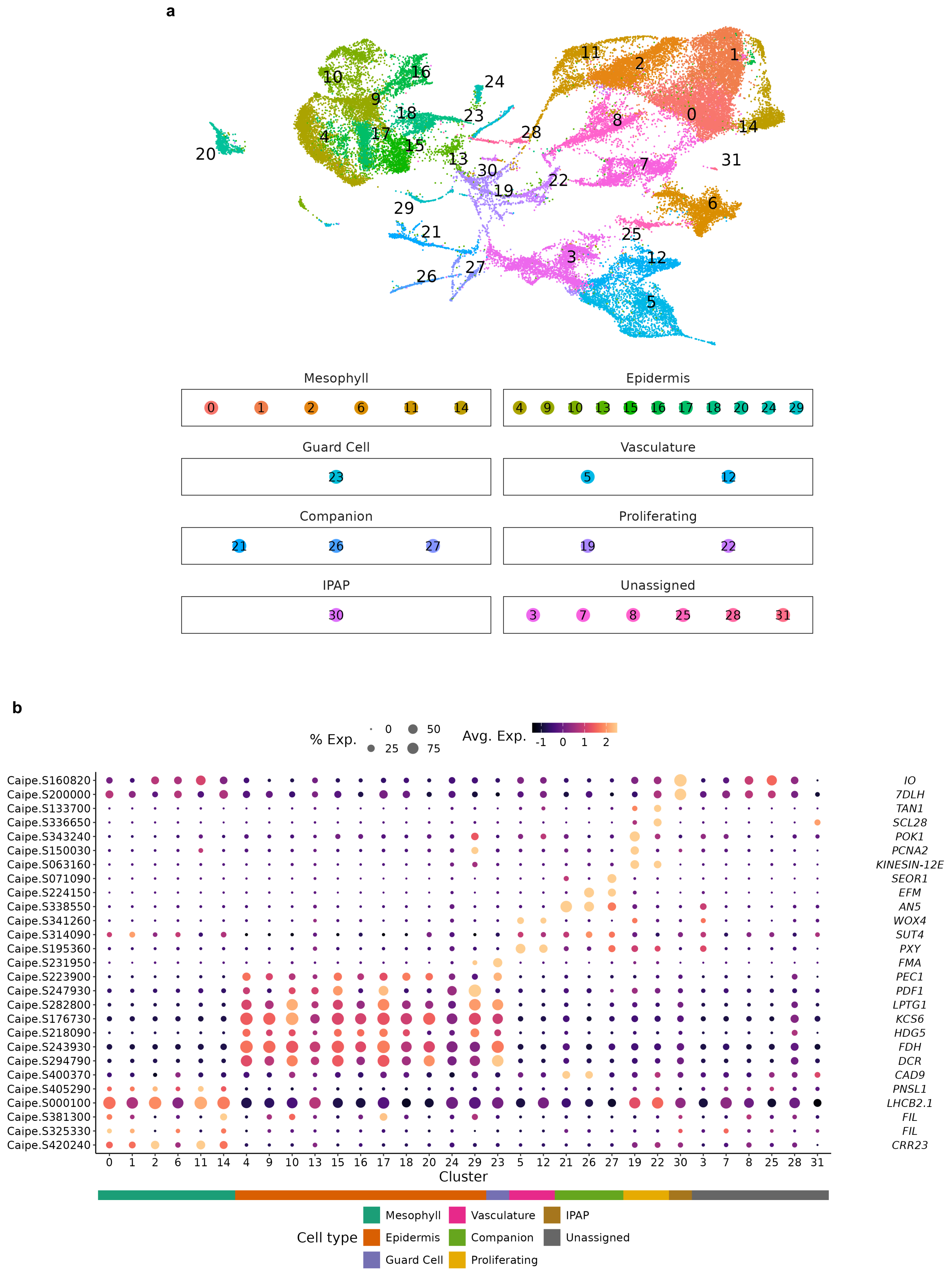

**Supplementary Fig. 4. Single nuclei RNA-seq *C. ipecacuanha* young leaves. a**, UMAP of average gene expression of two biological replicates of *C. ipecacuanha* young leaves ($n = 23{,}702$ cells for replicate 1, and 20,299 cells for replicate 2). Cell clusters were annotated as cell types using marker genes shown in b. IPAP, Internal Phloem Associated Parenchyma. **b**, Single-cell gene expression dot plot heatmap showing expression of orthologs of previously published marker genes for different cell types in other species (Supplementary Table 7). Color scale shows the average scaled expression of each gene over the different cell clusters. Dot sizes indicate the fraction of cells of each cell cluster in which a given gene is expressed.

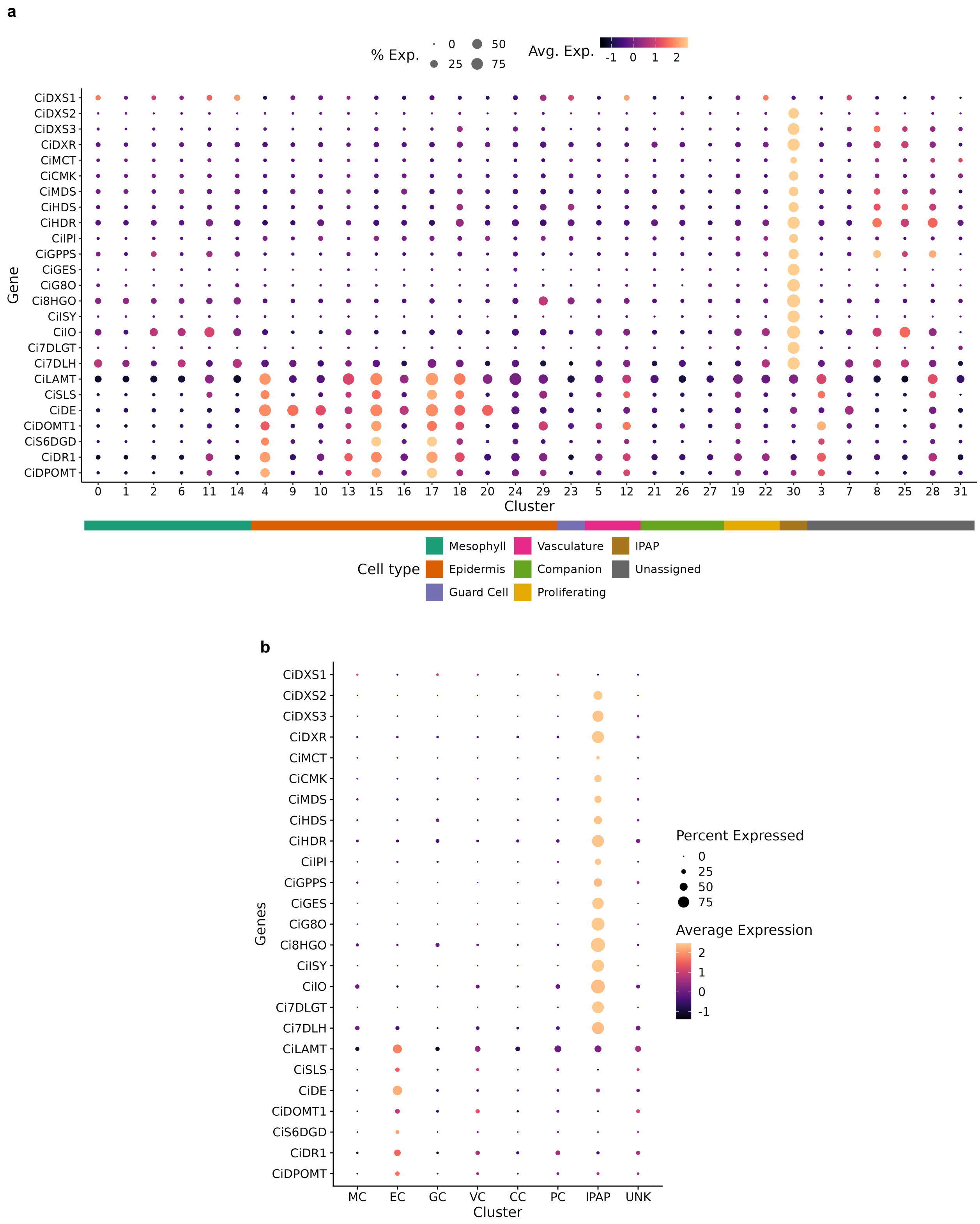

**Supplementary Fig. 5. Expression of iridoid and ipecac alkaloid pathway genes in single cell clusters.** The expression of orthologs of previously identified 2-C-methyl-D-erythritol 4-phosphate (MEP) pathway, secoiridoid pathway and downstream ipecac alkaloid pathway genes is shown (Colinas *et al.*, 2025; Li *et al*, 2023). Color scale shows the average scaled expression of each gene over the different cell clusters. Dot sizes indicate the fraction of cells of each cell cluster in which a given gene is expressed. **a**, Expression over 31 cell clusters. **b**, Expression over cell groups of identified cell types. MC, mesophyll cells; EC, epidermal cells; GC, guard cells; VC, vascular cells; CC, companion cells; PC, proliferating cells; IPAP, Internal Phloem Associated Parenchyma; UNK, unassigned.

**a**

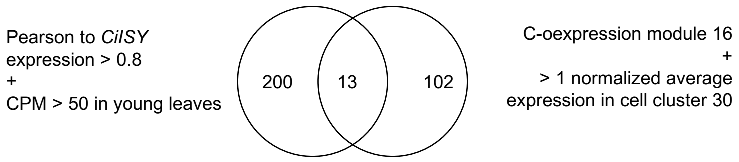

**List of 13 genes**

| GeneID | functional annotation | name / description | TPM in cell cluster 30 |
|---|---|---|---|
| Caipe.S388690 | alcohol dehydrogenase | Ci8HGO | 4.63 |
| Caipe.S160820 | cytochrome P450, family 76, subfamily C, polypeptide | CiIO | 4.27 |
| **Caipe.S125200** | **methyl esterase** | **CiICYC** | **3.99** |
| Caipe.S125190 | cytochrome P450, family 76, subfamily C, polypeptide | CiG8O | 3.71 |
| Caipe.S184100 | terpene synthase | CiGES | 3.64 |
| Caipe.S366190 | GroES-like zinc-binding dehydrogenase family protein | CYPADH ortholog | 3.45 |
| Caipe.S200000 | cytochrome P450, family 72, subfamily A, polypeptide | Ci7DLH | 3.39 |
| Caipe.S317530 | NAD(P)-binding Rossmann-fold superfamily protein | CiISY paralog | 3.33 |
| Caipe.S244560 | 1-deoxy-D-xylulose 5-phosphate reductoisomerase | DXR ortholog | 3.33 |
| Caipe.S232250 | Deoxyxylulose-5-phosphate synthase | DXS ortholog | 3.26 |
| Caipe.S317540 | NAD(P)-binding Rossmann-fold superfamily protein | CiISY | 3.04 |
| Caipe.S364920 | UDP-glucosyl transferase 85A3 | Ci7DLGT | 2.81 |
| Caipe.S351880 | sulfotransferase | | 2.57 |

**b**

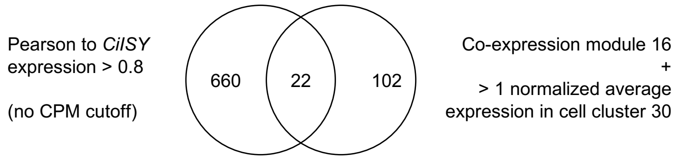

**List of 9 additional genes**

| GeneID | functional annotation | name / description | TPM in cell cluster 30 |
|---|---|---|---|
| Caipe.S317500 | F-BOX WITH WD-40 | | 3.81 |
| Caipe.S135190 | basic helix-loop-helix (bHLH) DNA-binding superfamily protein | BIS ortholog | 2.68 |
| Caipe.S429940 | Phosphatidylinositol-4-phosphate 5-kinase family protein | | 2.37 |
| Caipe.S215190 | Acyl-CoA N-acyltransferases (NAT) superfamily protein | | 2.11 |
| Caipe.S092330 | hypothetical protein | | 2.07 |
| Caipe.S135260 | basic helix-loop-helix (bHLH) DNA-binding superfamily protein | BIS ortholog | 1.82 |
| Caipe.S108310 | basic helix-loop-helix (bHLH) DNA-binding superfamily protein | | 1.42 |
| Caipe.S135200 | basic helix-loop-helix (bHLH) DNA-binding superfamily protein | BIS ortholog | 1.30 |
| Caipe.S352430 | Stress responsive alpha-beta barrel domain protein | | 1.11 |

**Supplementary Fig. 6. Combination of bulk and single nuclei RNA-seq co-expression analysis.** Because it was unclear which type of enzyme would catalyze this cyclization reaction, we attempted to narrow down the list of candidates through co-expression analyses rather than using functional annotation. To achieve the most highly resolved co-expression list, candidate genes from tissue specific co-expression analysis (Pearson correlation to *CiISY* expression > 0.8) and cell type specific co-expression analysis (co-expression module 16) were overlayed. **a**, In addition to co-expression, only genes with high absolute expression in bulk RNA-seq (> 50 CPM in young leaves from plant 1) and snRNA-seq (normalized average expression > 1 in cell cluster 30) were included. The overlay contained the 13 genes listed with functional annotations based on sequence homology. ICYC is shown in bold. Blue indicates orthologs of known iridoid pathway genes, green indicates orthologs of known MEP pathway genes. CYPADH has been previously associated with iridoid biosynthesis but its exact function could not be determined (Brown *et al*, 2015). **b**, Without a cutoff for absolute expression in bulk RNA-seq, the list of genes common to both datasets contained 9 additional genes, including orthologs of the *C. roseus* bHLH iridoid synthesis (BIS) transcription factors that are known to induce expression of IPAP specific iridoid pathway genes (Colinas *et al*, 2021; Van Moerkercke *et al*, 2016; Van Moerkercke *et al*, 2015). Blue indicates orthologs of *C. roseus* BIS. CPM, counts per million; TPM, transcript per million; GES, geraniol synthase; G8H, geraniol 8-hydroxylase; 8HGO, 8-hydroxygeraniol oxidase; ISY, iridoid synthase; ICYC, iridoid cyclase; IO, iridoid oxidase; 7DLGT, 7-deoxyloganetic acid glucosidase; 7DLH, 7-deoxyloganic acid hydroxylase.

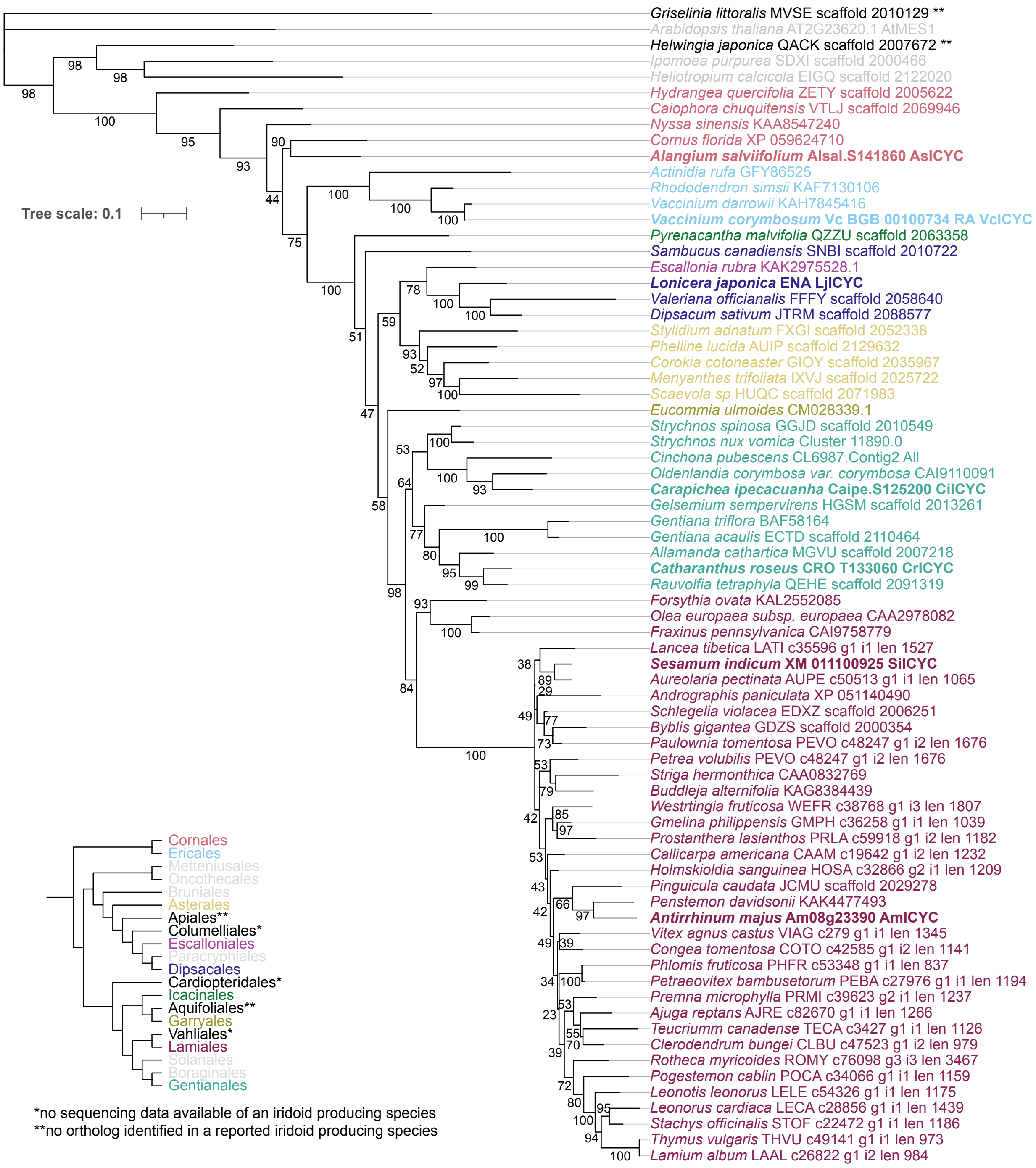

**Supplementary Fig. 7. Maximum-likelihood phylogenetic tree of ICYC amino acid sequences.** ICYC orthologs were identified in different asterid orders as indicated by the colour code. AtMES1 as well the closest ICYC homologs from non-iridoid producing species were included for comparison and do not cluster ICYCs. In the bottom left corner a tree showing all asterid orders is shown (Zuntini *et al*, 2024). An asterisk depicts that no sequencing data was publicly available from a reported iridoid producing species from the respective clade and thus ICYC presence could not be determined. Two asterisks depict that no ortholog could be found in the publicly available sequencing data a reported iridoid producing genus.

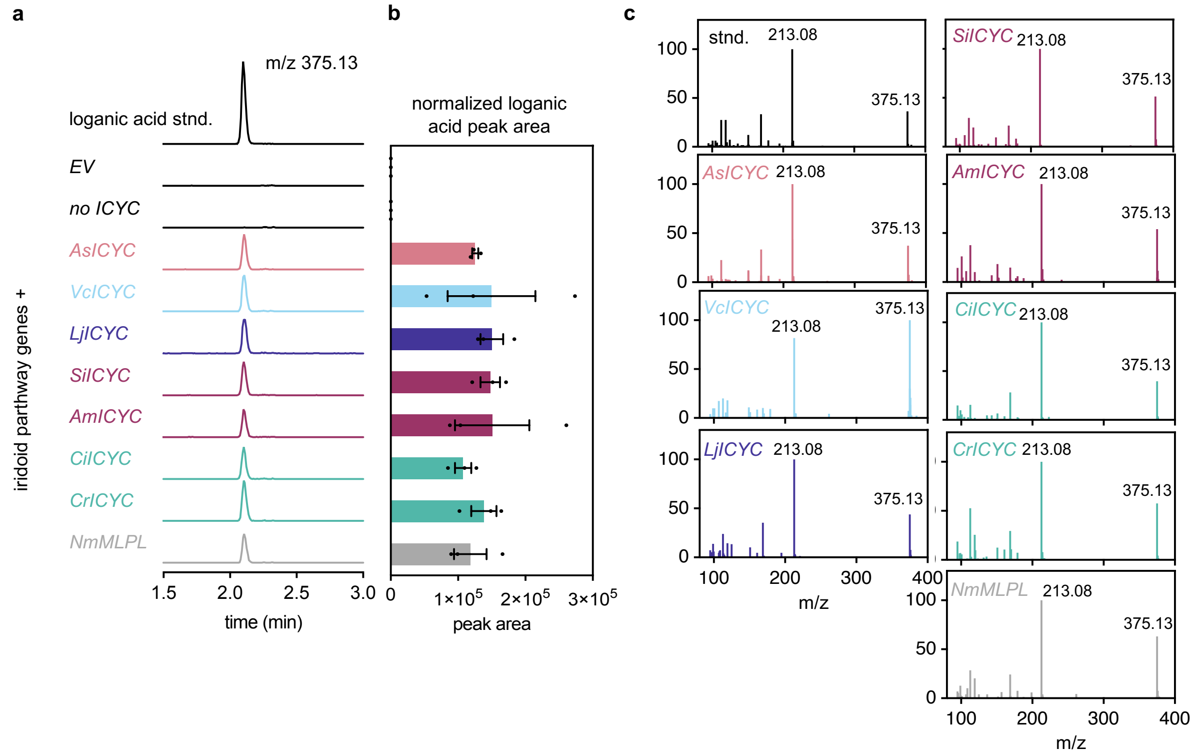

**Supplementary Fig. 8. ICYC orthologs from various Asterid orders enable loganic acid biosynthesis in *N. benthamiana*.** Full data corresponding to the experiment shown in Fig. 1e (main text). *C. ipecacuanha* loganic acid biosynthesis genes (see Fig. 1e for full list of genes) were co-overexpressed alongside *ICYC* orthologs. **a**, Extracted ion chromatograms (EICs) showing loganic acid *m/z* $[M\text{-}H]^-$ 375.13 of a representative biological replicate. **b**, LC-MS peak areas of loganic acid are shown as bars of the mean of three replicates, error bars are standard error of the mean. **c**, $MS^2$ fragmentation data for loganic acid peaks of different samples confirming peaks are identical to the commercial loganic acid standard. Stnd, standard.

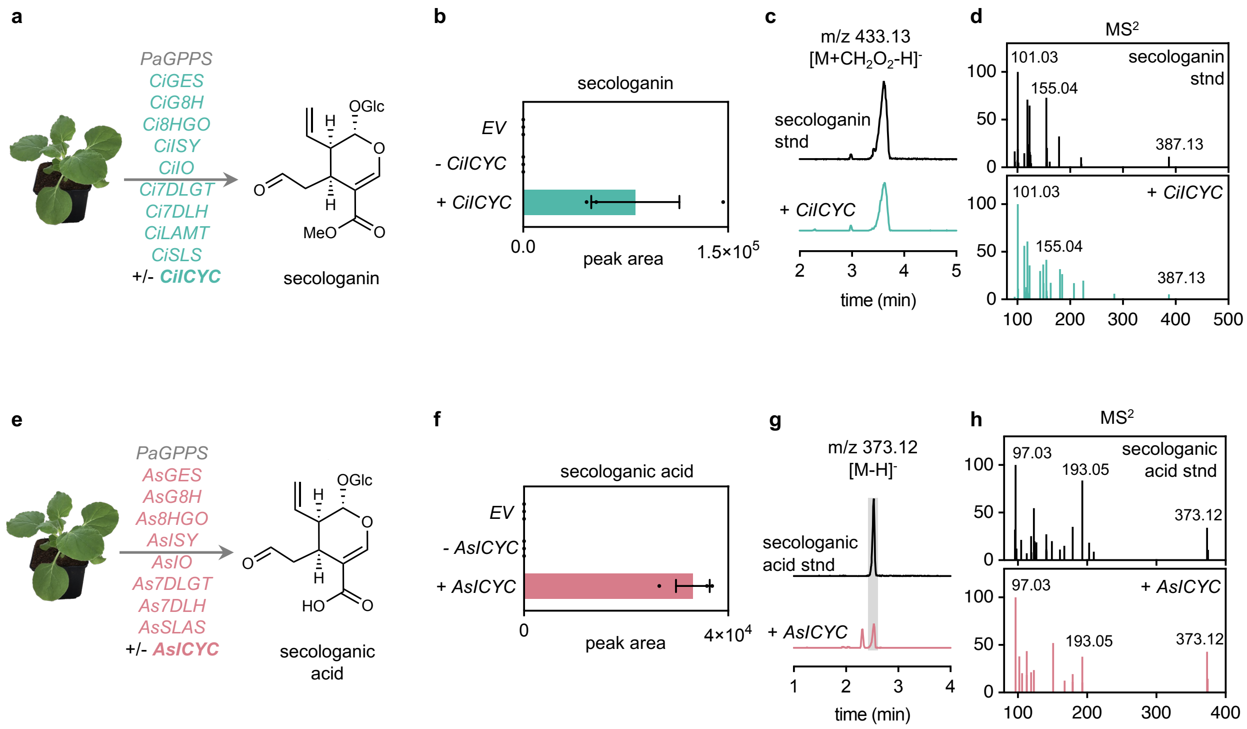

**Supplementary Fig. 9. Reconstitution of the secoiridoid pathway of *C. ipecacuanha* and *A. salviifolium* in *N. benthamiana*. a**-**d**, *C. ipecacuanha*. **e**-**h**, *A. salviifolium*. **a**, **e**, Genes overexpressed through agroinfiltration and expected products. **b**, **f**, Normalized peak areas for indicated products. LC-MS peak areas are shown as bars of the mean of three replicates, error bars are standard error of the mean. **c**, **g**, Extracted ion chromatograms alongside authentic standard. **d**, **h** $MS^2$ data of reconstitution product and standard confirming identity. EV, empty vector; *PaGPPS, Picea abies geraniol diphosphate synthase*; *GES, geraniol synthase; G8H, geraniol 8-hydroxylase; 8HGO, 8-hydroxygeraniol oxidase; ISY, iridoid synthase; ICYC, iridoid cyclase; IO, iridoid oxidase; 7DLGT, 7-deoxyloganetic acid glucosidase; 7DLH, 7-deoxyloganic acid hydroxylase; SLAS, secologanic acid synthase; LAMT, loganic acid methyltransferase; SLS, secologanin synthase; SLAS, secologanic acid synthase*.

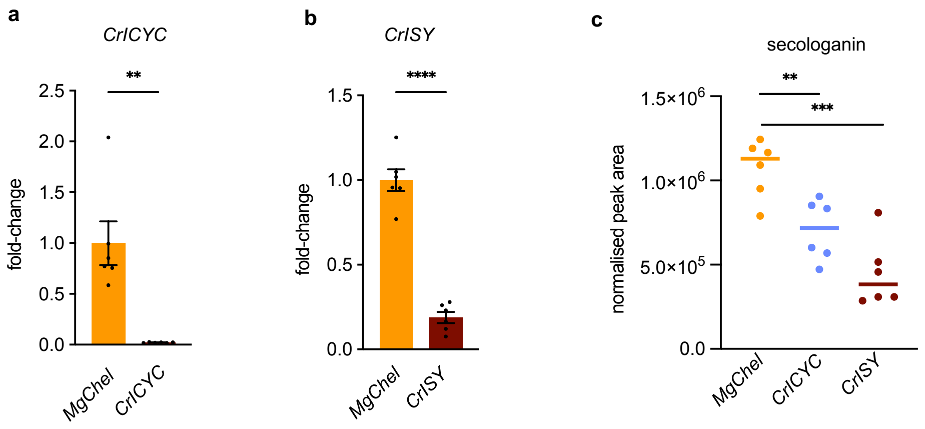

**Supplementary Fig. 10. Virus Induced Gene Silencing (VIGS) of *ICYC* and *ISY* in *C. roseus*.** *Magnesium Chelatase* (*MgChel*) was co-silenced in all cases to visualize silenced tissues. In the negative control (in orange) only *MgChel* was silenced. As a positive control *CrISY* was silenced. **a-b**, qPCR confirming the downregulation *CrICYC* (**a**) or *CrISY* (**b**) compared to *MgChel* negative control. Expression values are shown as fold-changes relative to expression in *MgChel* negative control. Bar graphs depict mean, error bars are standard error of the mean of N=6 biological replicates. Black dot symbols depict values for individual replicates. P-value of unpaired two-tailed ttest with Welch's test correction: **P=0.0061, ****P<0.0001. **c**, Secologanin levels in silenced plants compared to control. LC-MS peak areas normalized to internal standard are shown as individual dots of each replicate, the line depicts the mean of N=6 replicates. P-value of unpaired two-tailed ttest with Welch's test correction: **P=0.0047, ***P=0.0002.

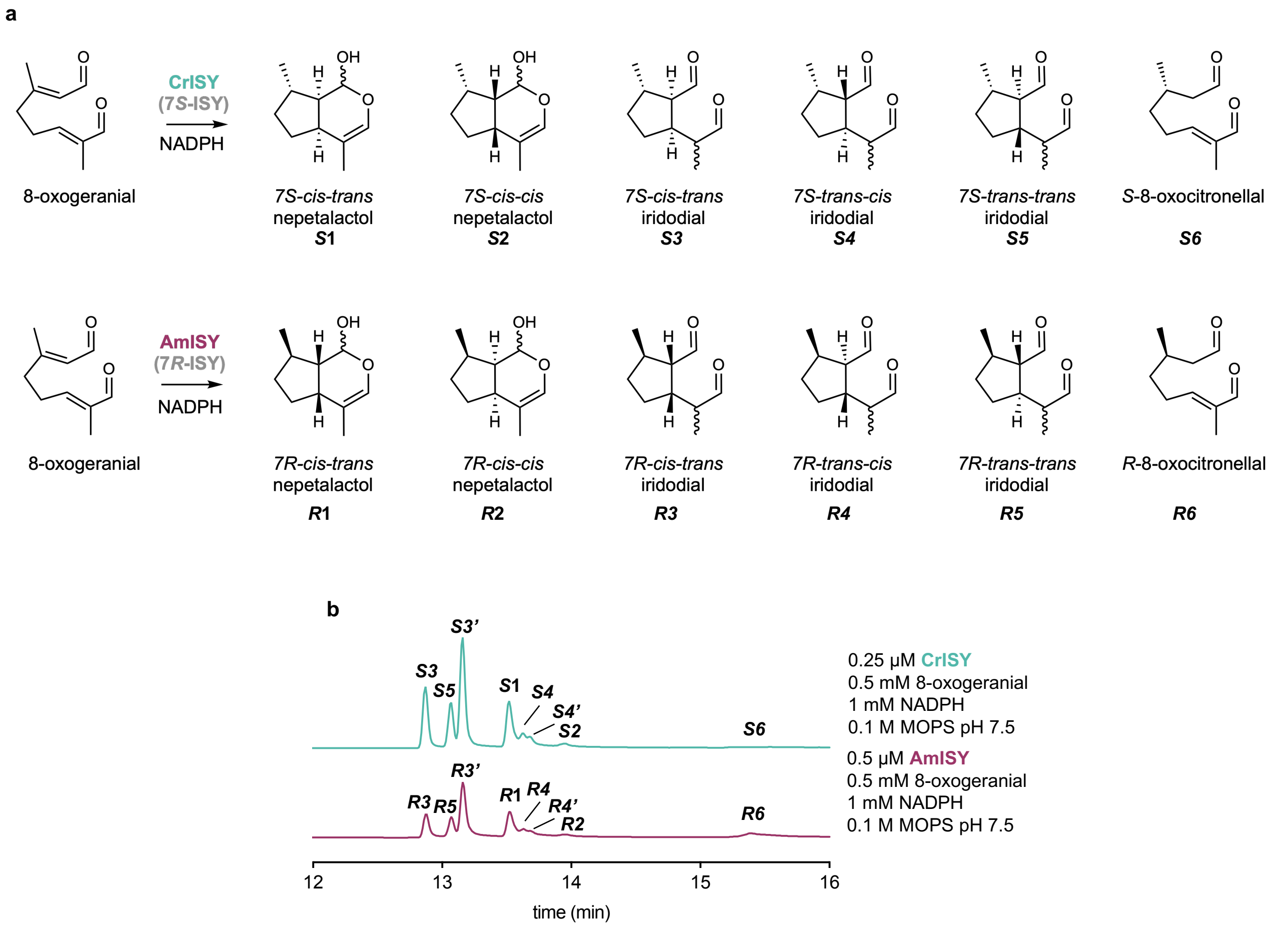

**Supplementary Fig. 11. Side products formed by CrISY and AmISY in the absence of cyclase.** **a**, Product structures formed by *C. roseus* iridoid synthase (CrISY). **b**, Product structures formed by *Antirrhinum majus* ISY (AmISY). **c**, GC-MS total ion chromatograms (TICs) of ISY catalyzed reactions as indicated. Note that enantiomers co-elute on an achiral column used here. Chromatograms of products were consistent with previously reported work and peak identification was inferred from the previously published profiles (Kries *et al*, 2017; Lichman *et al*, 2019).

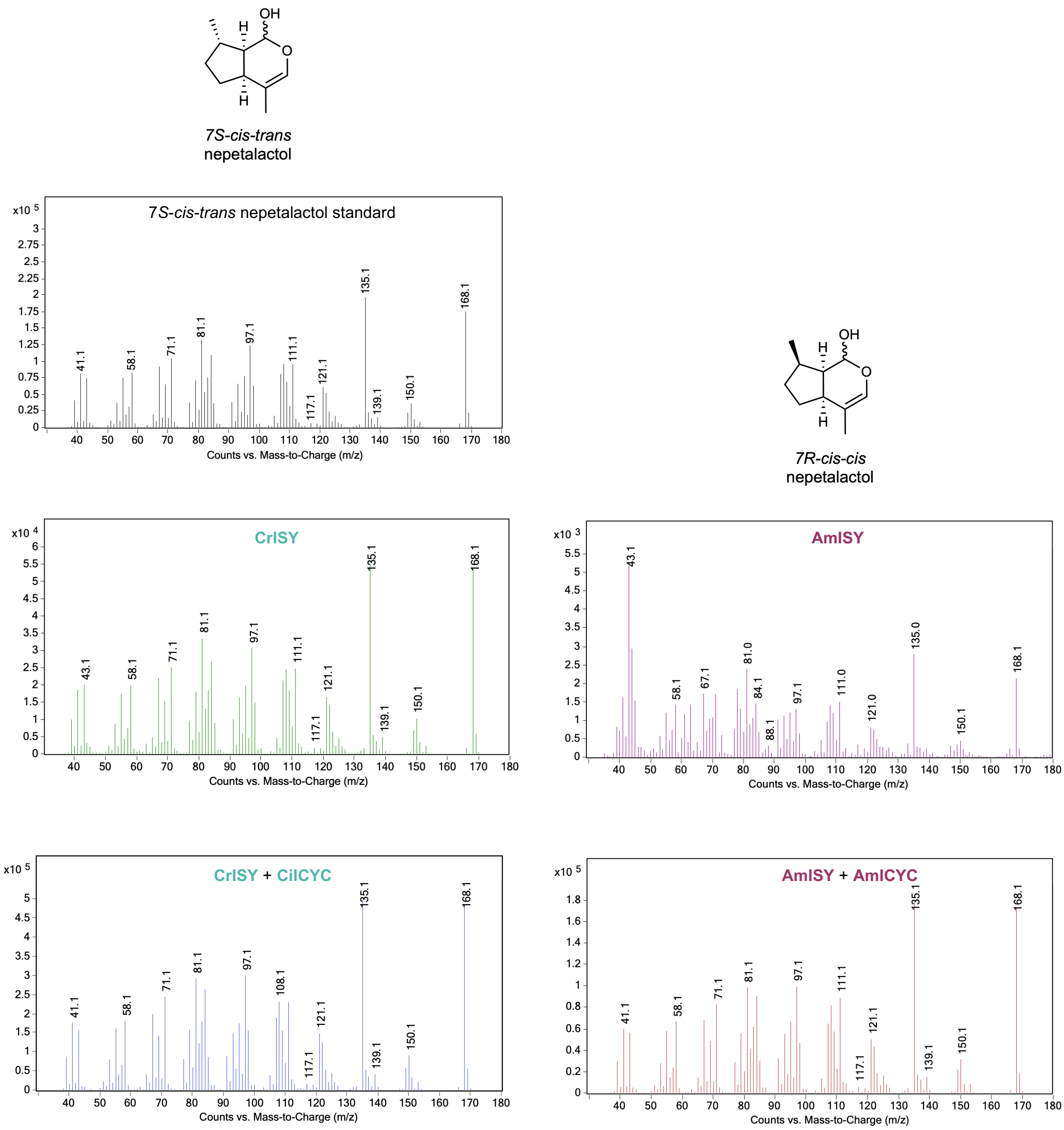

**Supplementary Fig. 12. Electron ionization (EI) spectra of nepetalactol standard and enzymatic products.** Spectra were obtained from the center of the peaks of chromatograms shown in Fig. 2 b, c.

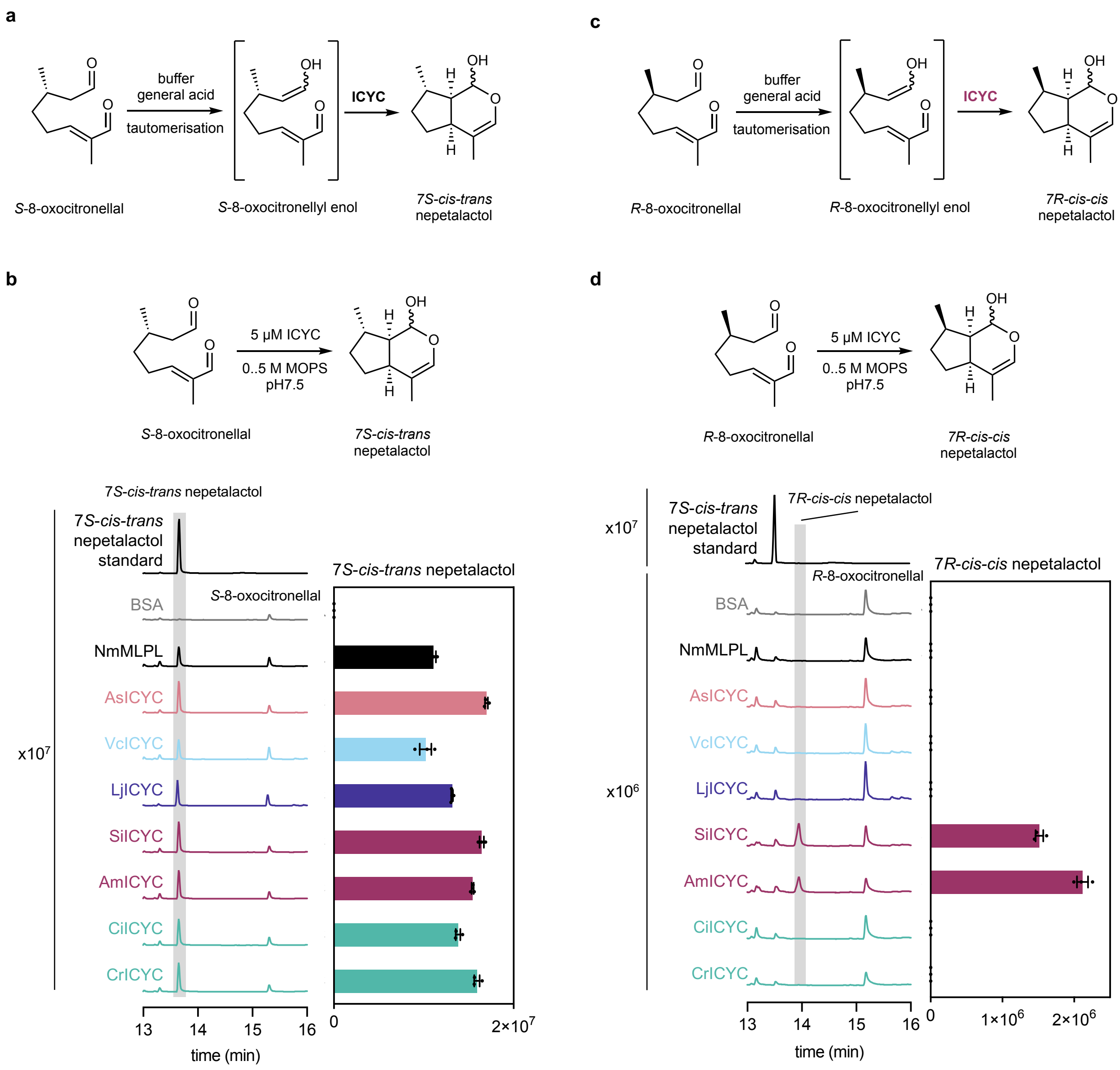

**Supplementary Fig. 13. ICYC activity with 8-oxocitronellal under tautomerization inducing conditions. a**, **c**, Under high general acid concentrations (0.5 M MOPS, pH 7.5) *S*-8-oxocitronellal (**a**) or *R*-8-oxocitronellal (**c**) partially tautomerize to the cyclase substrate 8-oxocitronellyl enol (Lichman *et al.*, 2019). **b**, Assays with *S*-8-oxocitronellal (0.5 mM) and ICYC or NmMLPL, or BSA as negative control for 16 hours. 7*S-cis-trans* nepetalactol was formed in similar amounts by all cyclases but not in the negative control **d**, Assays with *R*-8-oxocitronellal (0.1 mM due to limited substrate availability) for 16 hours. A peak identified as 7*R-cis-cis* nepetalactol (see Fig. 2d-f, main text) formed specifically in the presence of SiICYC and AmICYC but not with other ICYC orthologs. The results are consistent with the results observed for ISY-ICYC combined assays.

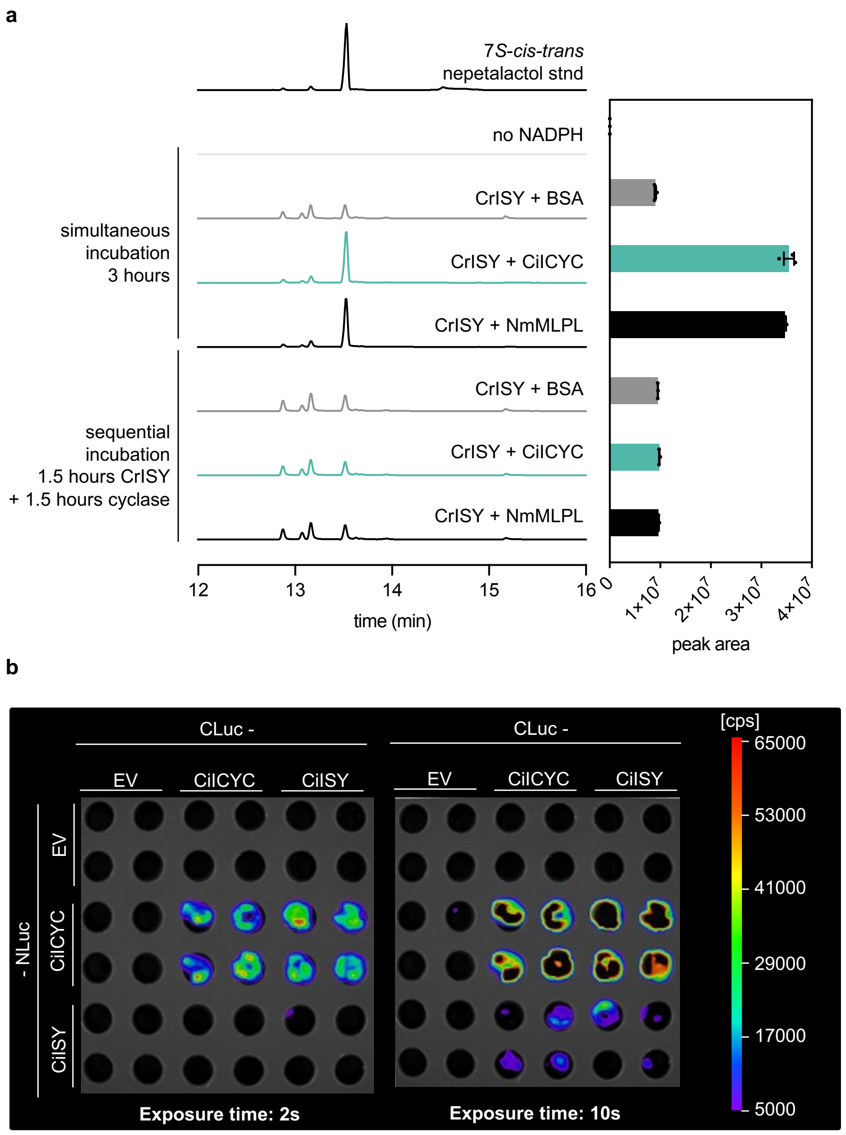

**Supplementary Fig. 14. ISY and ICYC must be simultaneously present for nepetalactol formation and may interact. a**, Enzyme assays under the conditions shown in Fig. 2a (main text) but with ICYC, NmMLPL or BSA added simultaneously for 3 hours or after 1.5 hours reaction with CrISY alone (sequential incubation). Interestingly, nepetalactol is only formed to higher amounts in simultaneous incubations. This could indicate that the formed ISY product 8-oxocitronellyl enol, which is the substrate for the cyclases, is unstable und must be immediately taken up by the cyclases. **b**, Split-Luciferase assay with CiICYC and CiISY in *N. benthamiana*. The C-terminal part of luciferase (CLuc) and the N-terminal part of luciferase (NLuc) are always fused N-terminally or C-terminally, respectively, to the target protein (Chen *et al*, 2008). Empty vector (EV) contained non-fused CLuc or NLuc, respectively, and served as negative controls. Leaf disks were cut from four biological replicates of agroinfiltrated *N. benthamiana* plants, transferred to a well plate, where luciferin was added, and imaged using a Nightshade camera (see methods for details). Images are pictures of the same leaf disks with different indicated exposure times. The assays indicate that CiICYC interacts with itself and that CiICYC interacts with CiISY. Note that when NLuc is fused to CiISY interaction is weaker and only revealed at longer exposure times.

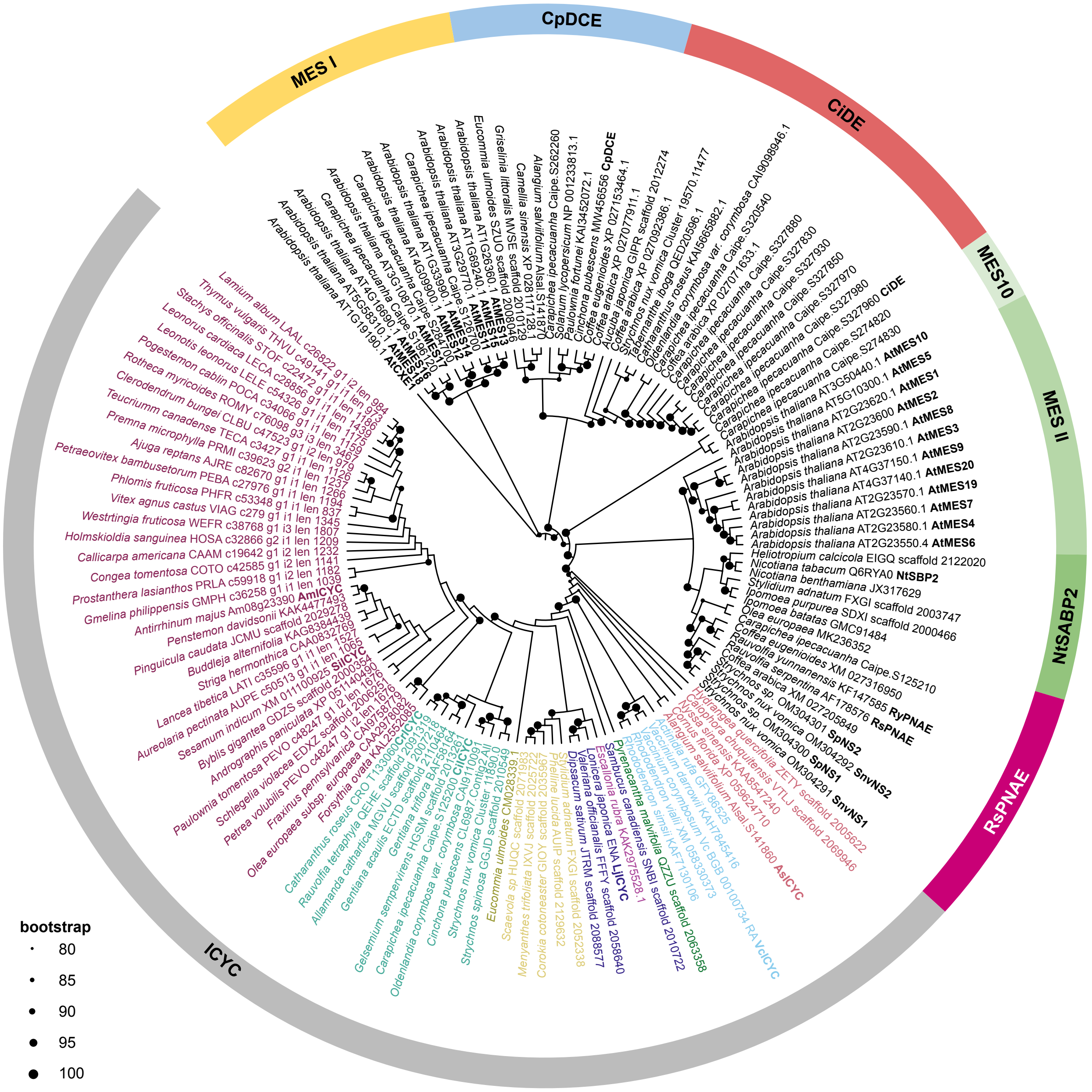

**Supplementary Fig. 15. Maximum likelihood tree of ICYC and methylesterase amino acid sequences.** *Arabidopsis thaliana* methylesterases (AtMES) were included as well as all *C. ipecacuanha* MES homologs for comparison. *A. thaliana* Carboxylesterase 1 (AtCXE)1 served as outgroup. *Nicotiana tabacum* salicylic acid binding protein 2 (SABP2) has been described as a homolog of AtMES' (Vlot *et al*, 2008). MES form distinct clades named I and II whereas MES10-like form a small separate clade. Different alkaloid esterases are found in well separated clades, each clade is named after a characterized member: *Cinchona pubescens* dihydrocorynantheine aldehyde esterase (CpDCE), *Carapichea ipecacuanha* deacetyl(iso)ipecoside esterase (CiDE), *Rauvolfia serpentina* polyneuridine aldehyde esterase (RsPNAE). Enzymes characterized in this or other studies are labelled with names in bold (Chaffin *et al*, 2024; Colinas *et al.*, 2025; Dogru *et al*, 2000; Hong *et al*, 2022; Trenti *et al*, 2021). NS, norfluorocurarine synthase.

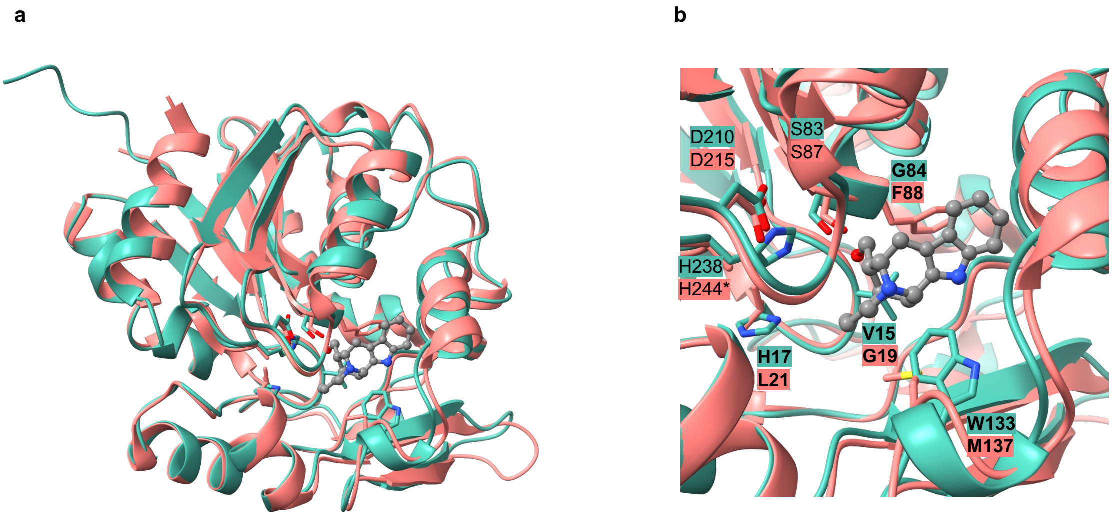

**Supplementary Fig. 16. Structural overlay of CiICYC model with RsPNAE structure**. Alphafold3 model of CiICYC (in green) was aligned with the crystal structure obtained for *Rauvolfia serpentina* polyneuridine aldehyde esterase (RsPNAE, in salmon) in complex with its product 16-epivellosimine (in grey) (PDB 3GZJ) (Yang *et al*, 2009). RsPNAE is the closest homolog of ICYC for which a crystal structure is available. **a**, Overview of structural alignment showing same protein folds. **b**, Closeup view of the active site. Amino acid residues highlighted in green and shown as sticks were chosen for mutation in CiICYC, corresponding amino acid residues in RsPNAE are also highlighted. In bold, amino acid residues that differed between the two proteins. Asterisk indicates that the native RsPNAE H244 was mutated to alanine in this protein crystal structure.

a b

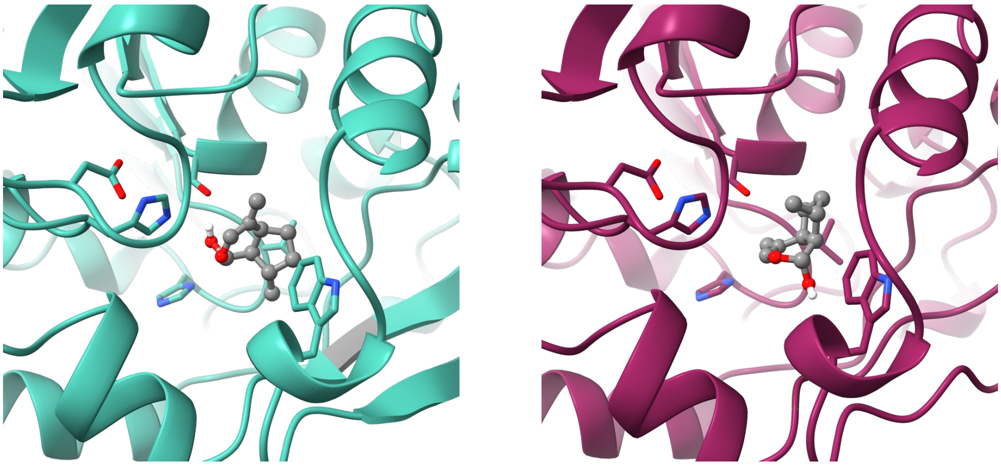

**Supplementary Fig. 17. Docking of nepetalactol stereoisomers to CiICYC and AmICYC models**. **a**, CiICYC alphafold3 model with 7*S*-*cis*-*trans* nepetalactol docked (AutoDock Vina) in active site. **b**, AmICYC alphafold3 model docked with 7*R*-*cis*-*cis* nepetalactol (Autodock Vina). Amino acid residues shown as sticks were targeted in CiICYC for mutation (Fig 3, main text) and are conserved in all ICYCs.

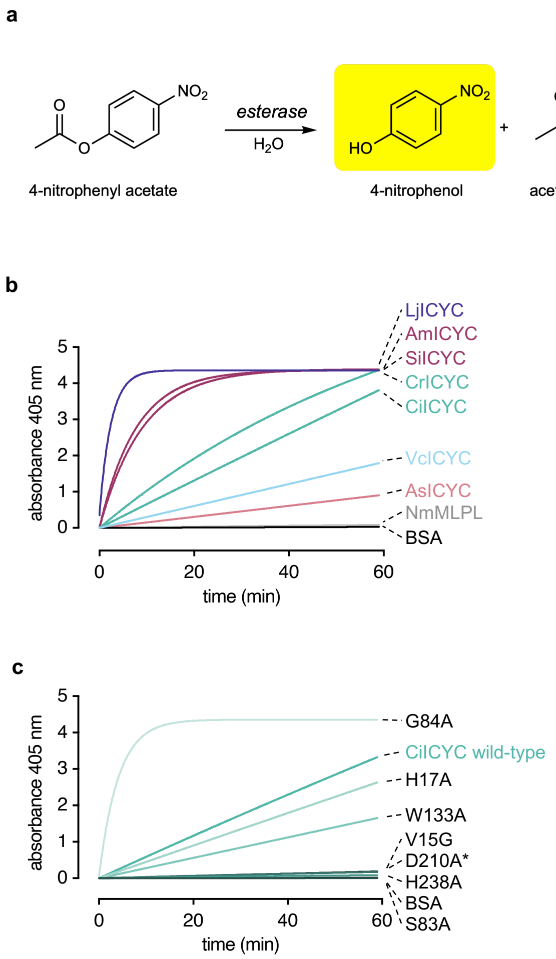

**Supplementary Fig. 18. Esterase activity of ICYC.** **a**, Reaction scheme of the assay. Esterase activity of the proteins were assessed with 4-nitrophenyl acetate as a substrate. The formation of product is assessed by measuring absorbance of the 4-nitrophenol (yellow) at 405 nm. **b**, Esterase activity assays of 0.5 µM ICYC orthologs compared to NmMLPL and the negative control BSA. Esterase activity greatly varies among ICYC proteins but is detectable for all ICYC orthologs. **c**, Esterase activity assays of 0.5 µM CiICYC wild-type and mutant proteins (see Fig. 3, main text). Mutations in the catalytic triad led to complete abolishment of esterase activity (the asterisk indicates that D210A mutant was poorly soluble, thus activity could not be assessed reliably). Esterase activity appeared to be poorly correlated with cyclase activity (Fig. 3, main text). Each curve was created by curve fitting of absorbance values of three replicates. Absorbance was measured every minute on a microplate reader (see methods for details).

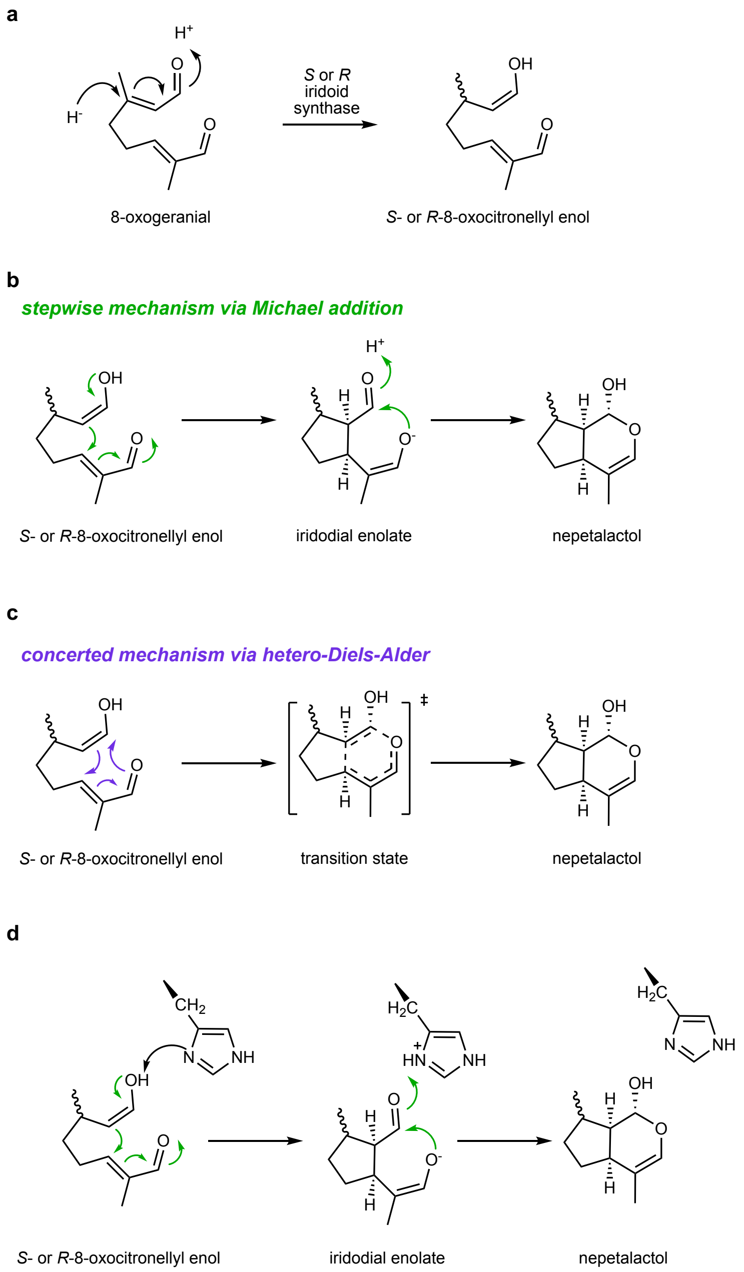

**Supplementary Fig. 19. Possible cyclization mechanisms. a**, 7*S*-iridoid synthase generates *S*-8-oxocitronellyl enol intermediate. **b**, Stepwise cyclization mechanism via Michael addition occurs via a five-membered ring intermediate. Previous studies using substrate analogs suggest that the spontaneous cyclization occurs via a stepwise Michael addition reaction mechanism (Lindner *et al*, 2014). **c**, A possible alternative mechanism is a concerted cyclization via hetero-Diels-Alder. **d**, Proposed role of the active site histidine (H238 in CiICYC) in a putative Michael addition type mechanism.

### Supplementary Tables (separate excel file)

**Supplementary Table 1. Sequencing data generated and used in this study.**
**Supplementary Table 2. Genome Assembly Metrics**
**Supplementary Table 3. Benchmarking universal single copy orthologs (BUSCO) results on the genome assemblies and annotation.**
**Supplementary Table 4. Repetitive Sequence Content**
**Supplementary Table 5. Gene Annotation Summary**
**Supplementary Table 6. Details of single nuclei RNA-seq libraries post processing**
**Supplementary Table 7. Marker genes used to assign cell types**
**Supplementary Table 8. List of primers used in this study.**
**Supplementary Table 9. Accession numbers of genes described in this study.**
**Supplementary Table 10. Sequences used to construct phylogenetic trees.**

### Supplementary Dataset 1. Cell cluster co-expression modules (separate excel file)

### Supplementary references

Brown S, Clastre M, Courdavault V, O'Connor SE (2015) De novo production of the plant-derived alkaloid strictosidine in yeast. *Proc Natl Acad Sci U S A* 112: 3205-3210

Chaffin TA, Wang W, Chen J-G, Chen F (2024) Function and Evolution of the Plant MES Family of Methylesterases. *Plants* 13

Chen H, Zou Y, Shang Y, Lin H, Wang Y, Cai R, Tang X, Zhou JM (2008) Firefly luciferase complementation imaging assay for protein-protein interactions in plants. *Plant Physiol* 146: 368-376

Colinas M, Morweiser C, Dittberner O, Chioca B, Alam R, Leucke H, Nakamura Y, Serna Guerrero DA, Heinicke S, Kunert M *et al* (2025) Ipecac alkaloid biosynthesis in two evolutionarily distant plants. *Nat Chem Biol*

Colinas M, Pollier J, Vaneechoutte D, Malat DG, Schweizer F, De Milde L, De Clercq R, Guedes JG, Martinez-Cortes T, Molina-Hidalgo FJ *et al* (2021) Subfunctionalization of Paralog Transcription Factors Contributes to Regulation of Alkaloid Pathway Branch Choice in Catharanthus roseus. *Front Plant Sci* 12: 687406

Dogru E, Warzecha H, Seibel F, Haebel S, Lottspeich F, Stockigt J (2000) The gene encoding polyneuridine aldehyde esterase of monoterpenoid indole alkaloid biosynthesis in plants is an ortholog of the alpha/betahydrolase super family. *Eur J Biochem* 267: 1397-1406

Hong B, Grzech D, Caputi L, Sonawane P, Lopez CER, Kamileen MO, Hernandez Lozada NJ, Grabe V, O'Connor SE (2022) Biosynthesis of strychnine. *Nature* 607: 617-622

Kang M, Fu R, Zhang P, Lou S, Yang X, Chen Y, Ma T, Zhang Y, Xi Z, Liu J (2021) A chromosome-level Camptotheca acuminata genome assembly provides insights into the evolutionary origin of camptothecin biosynthesis. *Nat Commun* 12: 3531

Kries H, Kellner F, Kamileen MO, O'Connor SE (2017) Inverted stereocontrol of iridoid synthase in snapdragon. *J Biol Chem* 292: 14659-14667

Li C, Wood JC, Vu AH, Hamilton JP, Rodriguez Lopez CE, Payne RME, Serna Guerrero DA, Gase K, Yamamoto K, Vaillancourt B *et al* (2023) Single-cell multi-omics in the medicinal plant Catharanthus roseus. *Nat Chem Biol* 19: 1031-1041

Lichman BR, Kamileen MO, Titchiner GR, Saalbach G, Stevenson CEM, Lawson DM, O'Connor SE (2019) Uncoupled activation and cyclization in catmint reductive terpenoid biosynthesis. *Nat Chem Biol* 15: 71-79

Lindner S, Geu-Flores F, Brase S, Sherden NH, O'Connor SE (2014) Conversion of substrate analogs suggests a Michael cyclization in iridoid biosynthesis. *Chem Biol* 21: 1452-1456

Miller JC, Hollatz AJ, Schuler MA (2021) P450 variations bifurcate the early terpene indole alkaloid pathway in Catharanthus roseus and Camptotheca acuminata. *Phytochemistry* 183: 112626

Trenti F, Yamamoto K, Hong B, Paetz C, Nakamura Y, O'Connor SE (2021) Early and Late Steps of Quinine Biosynthesis. *Org Lett* 23: 1793-1797

Van Moerkercke A, Steensma P, Gariboldi I, Espoz J, Purnama PC, Schweizer F, Miettinen K, Vanden Bossche R, De Clercq R, Memelink J *et al* (2016) The basic helix-loop-helix transcription factor BIS2 is essential for monoterpenoid indole alkaloid production in the medicinal plant Catharanthus roseus. *Plant J* 88: 3-12

Van Moerkercke A, Steensma P, Schweizer F, Pollier J, Gariboldi I, Payne R, Vanden Bossche R, Miettinen K, Espoz J, Purnama PC *et al* (2015) The bHLH transcription factor BIS1 controls the iridoid branch of the monoterpenoid indole alkaloid pathway in <i>Catharanthus roseus</i>. *Proceedings of the National Academy of Sciences* 112: 8130-8135

Vlot AC, Liu PP, Cameron RK, Park SW, Yang Y, Kumar D, Zhou F, Padukkavidana T, Gustafsson C, Pichersky E *et al* (2008) Identification of likely orthologs of tobacco salicylic acid-binding protein 2 and their role in systemic acquired resistance in Arabidopsis thaliana. *Plant J* 56: 445-456

Yang L, Hill M, Wang M, Panjikar S, Stockigt J (2009) Structural basis and enzymatic mechanism of the biosynthesis of C9- from C10-monoterpenoid indole alkaloids. *Angew Chem Int Ed Engl* 48: 5211-5213

Zuntini AR, Carruthers T, Maurin O, Bailey PC, Leempoel K, Brewer GE, Epitawalage N, Françoso E, Gallego-Paramo B, McGinnie C *et al* (2024) Phylogenomics and the rise of the angiosperms. *Nature*
